# SEA CDM: Study-Experiment-Assay Common Data Model and Databases for Cross-Domain Data Integration and Analysis

**DOI:** 10.1101/2025.08.26.671804

**Authors:** Anthony Huffman, Feng-Yu Yeh, Junguk Hur, Jie Zheng, Anna Maria Masci, Guanming Wu, Cui Tao, Brian Athey, Yongqun He

## Abstract

With the increasing volume of biomedical experimental data, standardizing, sharing, and integrating heterogeneous experimental data across domains has become a major challenge. To address this challenge, we have developed an ontology-supported Study-Experiment-Assay (SEA) common data model (CDM), which includes 10 core and 3 auxiliary classes based on object-oriented modeling. SEA CDM uses interoperable ontologies for data standardization and knowledge inference. Building on the SEA CDM, we developed the Ontology-based SEA Network (OSEAN) relational database and knowledge graph, along with a set of ETL (Extract, Transform, Load) and query tools, and further applied them to represent 1,278 immune studies with over two million samples from three resources: VIGET, ImmPort, and CELLxGENE. Using simple, robust queries and analyses, our research identified multiple scientific insights into sex-specific immune responses, such as neutrophil degranulation and TNF binding to physiological receptors, following live attenuated and trivalent inactivated influenza vaccination. The novel SEA CDM system lays a foundation for establishing an integrative biodata ecosystem across biological and biomedical domains.

## Background

With intensive biological and biomedical research conducted, a large number of databases and resources have been developed to store increasingly complex data with various variables. For example, the ImmPort database is the world’s largest repository of public-domain immune response study data and analysis portal^1^. ImmPort stores the metadata of various immune studies. While ImmPort stores some processed data, it does not store specific high-throughput gene expression data, which is instead stored in the Gene Expression Omnibus (GEO) repository^2^. The Vaccine Immune Gene Expression Tool (VIGET)^3^ downloaded vaccine-related immune response metadata from ImmPort, processes gene expression data from GEO, and provides a user-friendly web interface for automated querying, processing, and statistical analysis of various metadata and gene expression data across different vaccine studies. The VIGET web interface is within the comprehensive Vaccine Investigation and Online Information Network (VIOLIN) vaccine database and analysis system^4^. The Chan Zuckerberg CELLxGENE Discover (CZ CELLxGENE) database stores many cell-level gene expression data such as single-cell or single-nucleus RNA-seq data^5^. Examples of other biomedical data resources include the Cancer Genome Atlas project for cancer (TCGA)^6^, the Kidney Precision Medicine Project for kidneys (KPMP)^7^, and the Human BioMolecular Atlas Program (HuBMAP) for human cells^8^.

It has been a major challenge to integrate heterogeneous data from different data resources. Different data resources tend to focus on specific domains, and the study types and variables collected in these resources often differ a lot, although overlaps do exist. While these databases individually may have specific guidelines designed to consistently report data within their domain, the guidelines are usually not generalizable to other databases. Integration is key to data being FAIR: Findable, Accessible, Interoperable, and Reusable^9^. The National Institutes of Health (NIH) promotes the data ecosystem via the Common Fund Data Ecosystem and individual NIH institutes (e.g., National Institute of Allergy and Infectious Diseases (NIAID)^10^. It is critical to address this issue at a fundamental level.

Ontology has been used as a key method for data standardization. An ontology, for biomedical research, is a hierarchical open-world knowledge graph about some real-world domain^11^. ImmPort uses ontologies such as the Vaccine Ontology (VO)^12^ and Ontology of Biomedical Investigation (OBI)^13^ for standardizing key concepts found within its domain, such as different diseases or assays used for immune studies. Ontologies provide a consistent way to name concepts and data and serve as a bedrock for data FAIRness. However, given different systems all using ontologies, the use of ontologies is a prerequisite rather than a sufficient condition.

There have been several attempts to create a data representation framework beyond a single domain. The Investigation-Study-Assay (ISA) framework^14^ was developed to provide a general framework for scientific investigations. The ISA-Tab tools were developed to aid in recording files into the ISA format^15^. Similarly, the Observational Medical Outcomes Partnership (OMOP) Common Data Model (CDM) is used for human electronic health records (EHR)^16^. OMOP CDM incorporates 39 distinct classes that can be leveraged for further health-related studies^16^. OMOP has been adopted by the All of Us Research Program^17^, the National Clinical Cohort Collaborative (N3C)^18^, and the Observational Health Data Sciences and Informatics (OHDSI)^19^ as part of a new standard. As such, while there is a framework for investigations and clinical visits, there is no framework that explicitly focuses on the integration of all biological experiments. This represents an obvious gap for furthering biomedical research.

In this manuscript, we report our creation and application of a Study-Experiment-Assay (SEA) CDM to better model studies within CELLxGENE, ImmPort, and our own VIGET^3^, which uses data from GEO and ImmPort. The consistent use of a SEA model facilitates integration across multiple data models, complementing ISA. As a use case, we represent vaccine immune study data from CELLxGENE and ImmPort to illustrate how SEA CDM works.

## Results

### Systematic SEA CDM for unified cross-domain biomedical study integration

**Figure 1** represents the overview of the SEA CDM, which contains 10 core classes and 3 accessory (or utility) classes based on object-oriented modeling. The class level representation of SEA CDM is implemented in two types of databases: a relational database (as seen in the OMOP CDM^16^), and a knowledge graph (KG) (as implemented in Neo4j). In the relational database setting, a class equals a table^20^, and in the KG setting, a class equals a node type^21^. Note that the classes in **Figure 1** are defined based on object-oriented modeling. Meanwhile, these classes can be classified here as material entity (blue), processes (red), and data (or information artifact, yellow) based on the Basic Formal Ontology^22^ and Information Artifact Ontology^23^. Such ontological classification better defines these classes.

**Figure 1.**
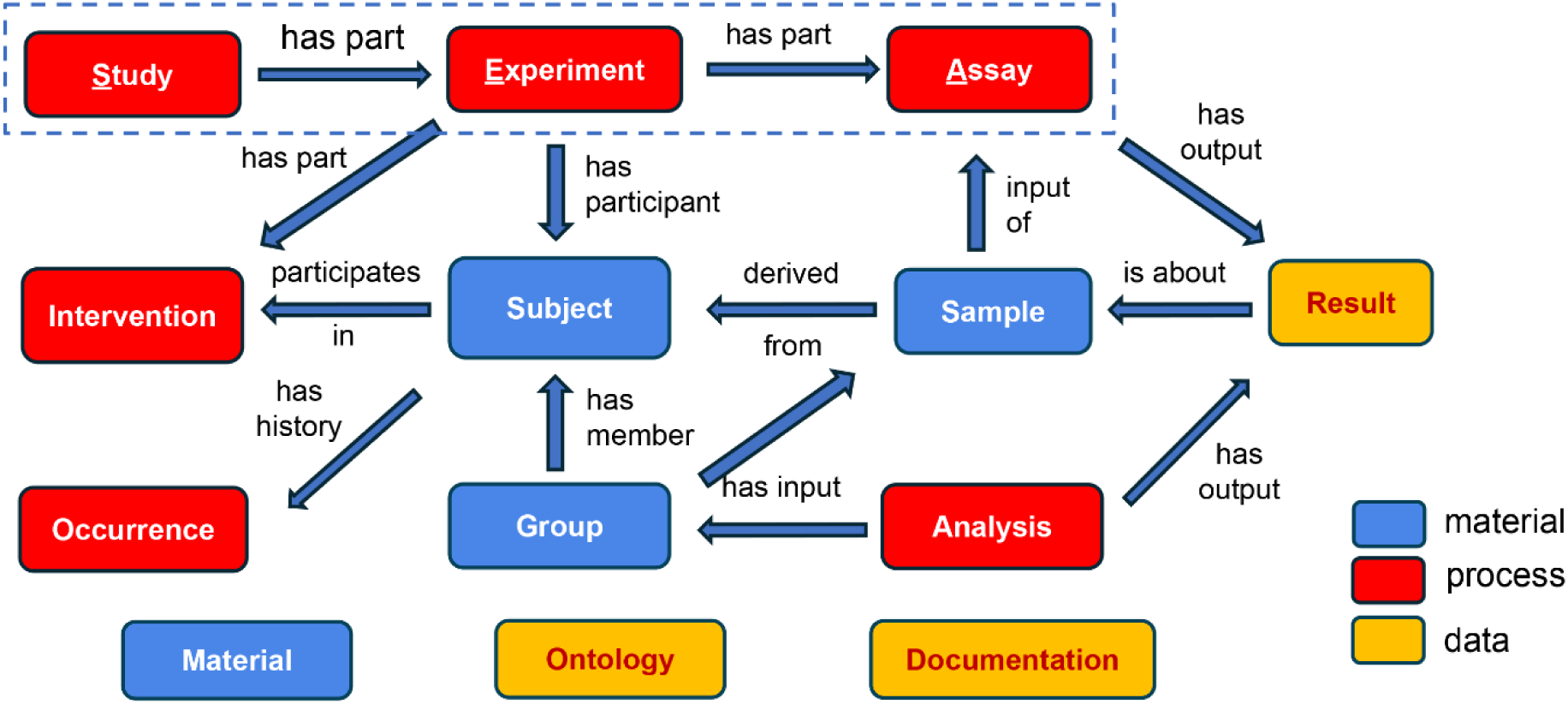
Overview of the “Study-Experiment-Assay” (SEA) common data model (CDM). This SEA CDM contains 13 classes, with 10 core classes linked by specific relations. Boxes in red represent classes for modeling processes, boxes in blue for material entities, and boxes in orange for data items. Classes represented by three floating boxes are accessory classes that help the core classes in data representation.

The 10 core SEA CDM classes include three high-level process-level classes: Study, Experiment, and Assay, which represent three interconnected processes in experimental studies. These classes are closely associated with the other core classes including three material entity level classes (i.e., *Subject*, *Sample*, and *Group*), three other process level classes (i.e., *Intervention*, *Occurrence*, and *Analysis*), and one data level class (i.e., *Result*). The basic logic here is that a study includes one or more experiments that generate samples, and one experiment contains one or more assays that utilize any samples generated by the experiment to generate specific results. Here, we focus on biomedical studies involving organisms, which can be humans or other animals (e.g., mice). The organisms may be intervened by some human-caused intervention procedure such as vaccination, drug treatment, device exposure, or surgery. The organisms can be grouped into different experimental groups, which can be used for statistical analysis. In addition to human-caused interventions, organisms may experience natural occurrences such as a history of some disease (e.g., diabetes) or an adverse event after intervention.

Many of these core SEA CDM classes are defined in a broad rather than a narrow sense. The *Study* class contains all investigations and studies related to addressing a scientific question or hypothesis. The *Experiment* class serves as a central node that connects the different aspects of a study and covers two types of ‘experiments’: human clinical visits and non-human animal model experiments. The *Assay* class covers both traditional assays generated from a sample and observations collected from a specimen. The *Sample* class also distinguishes between a sample that is collected from a subject (biosample, e.g., blood sample, urine sample) and one that has been obtained by processing biosample and used for later experimental investigation (expsample, e.g., RNA extract, kidney cell extract). Finally, the *Result* class of an assay can include the raw output of an assay, the transformed data of an assay, or the statistical analysis performed on the assay.

The 13 SEA CDM classes also include 3 accessory classes: *Material*, *Ontology*, and *Documentation*. The *Material* class includes the agents used in the experiments, such as a vaccine or saline, and the detailed agent information. The *Ontology* class covers the information of specific ontology (or ontologies) used in the study representation. The *Documentation* class includes the metadata and paths to specific documents, e.g., protocol, data files, or publications, and it contains specific information for the different processes found in SEA CDM. Further explanation on the SEA CDM model can be found on our website at (https://SEACDM.github.io/SEA-CDM/index.html).

**Table 1** lists representative class attributes of 6 out of 13 distinct classes in the SEA CDM. For example, SEA CDM *Subject’s* basic attributes include age, sex, species, human-specific race and ethnicity, and nonhuman strain. For a particular *Intervention* (e.g., vaccination), SEA CDM includes the material, intervention time, dose, route, and intervention type (e.g., vaccination, treatment, immunization). *Sample* attributes include collection method, collection time, biosample source, and experimental sample type. *Assay* attributes include assay type, organism inclusion (or not), reagents, and platform. *Result* attributes include datatype, original assay type, and file type. Note that when these object-oriented modeling classes and attributes are used for relational database development, the classes become the database tables, and the attributes become the columns of the database tables. Furthermore, the database tables may also include foreign keys as new columns to cross-link between tables (**Supplemental Table 1**).

**Table 1.** Representative classes and attributes (or properties) on SEA CDM. Each attribute can be used as a variable for scientific analysis.

| **Classes** | **Representative Attributes** |
| --- | --- |
| Subject | age, sex, species, race, ethnicity, strain |
| Intervention | intervention_type, material, intervention_time, dose, route |
| Sample | collection, collection_time, biosample_source, expsample_type |
| Assay | assay_type, organism_inclusion, reagents, platform |
| Result | datatype, original_assay_type, filetype |

The use of standardized ontologies is critical to the SEA CDM system. First, ontologies are used to standardize classes and attributes within SEA CDM. Each attribute within SEA CDM either represents metadata used to access or load in a file, or serves as a variable in a biological experiment. Here we use the term “variable” in a way that is aligned with the OBI^13^, where a variable is ‘A directive information entity that is about a data item which is realized through statistical analysis.” Each variable is paired with an ontology term that can be used to query for additional information. A set of reference interoperable ontologies was used to identify where to map attributes and variables from other programs to SEA CDM categories (See Methods).

Furthermore, ontologies provide additional information beyond the term names and identifiers. For example, given a vaccine name ‘Fluzone’ and its Vaccine Ontology (VO) identifier (VO_0000047), we can search VO and find more information about Fluzone, such as its classification as a trivalent inactivated influenza vaccine against influenza type A and B viral infection and its production by Sanofi Pasteur Limited^12^. In addition, ontology contains information about related terms and allows users to infer semantic relations among terms. **Figure 2A** shows an example of the hierarchy of ‘trivalent influenza vaccine’ defined in VO. Specific vaccines like Fluarix, Fluvirin, and Fluzone are defined as trivalent influenza vaccines. We can further query the ontology using tools such as a SPARQL query to retrieve related entities. For example, a simple SPARQL query script in **Figure 2B** would quickly identify 771 specific vaccines under the hierarchical category of ‘trivalent influenza vaccine’ in VO. Such a feature would support advanced semantic queries.

**Figure 2.**
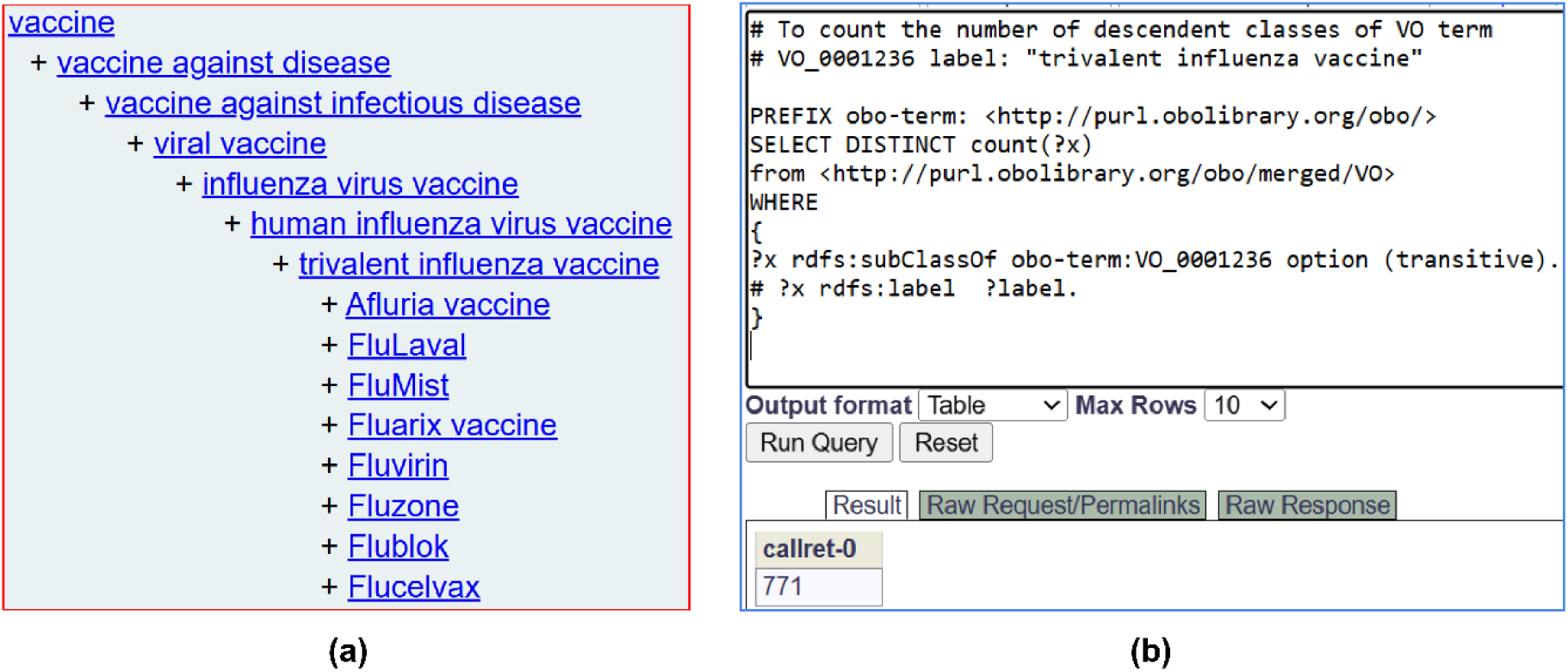
Ontological representation of key terms and their hierarchies. (A) a Vaccine Ontology (VO) hierarchy of trivalent influenza vaccine. (B) A SPARQL query for the total number of ‘trivalent influenza vaccine’ (VO_0001236) in VO.

### SEA CDM-based OSEAN database and knowledge graph generation

To better use the SEA CDM, we have developed a SEA CDM-based relational database called the Ontology-based SEA Network (OSEAN) database (OSEAN-DB) and a SEA CDM-based knowledge graph called OSEAN-KG. Both OSEAN-DB and OSEAN-KG use the same templates of data and follow the same SEA CDM format. **Supplemental File 1** contains all the SEA CDM templates and instructions for generating different tables (for the relational database) or classes (for knowledge graph). Each SEA CDM template file (.csv) contains a header row that lists class metadata and attributes. New entries are to be filled within a new row. Each attribute is by default paired with a column for its corresponding ontology ID from a reference ontology. Each csv file can be filled out manually or through automated software programs.

The schema for loading the MySQL OSEAN-DB is available on the GitHub website (https://sea-cdm.github.io/SEA-CDM/) and a Zenodo repository (https://zenodo.org/records/17770032) as detailed more in the section of Methods. The above newly converted SEA CDM tables can be used as inputs to develop the OSEAN-DB. Instructions to access a dump file containing the files needed to load OSEAN-DB or OSEAN-KG can be found as part of **Supplemental File 2**. Furthermore, we also provide instructions to load files from VIGET, ImmPort, CELLxGENE into SEA-CDM format as our use cases. More details are provided in the Methods section.

Below we will describe how the SEA CDM and its associated OSEAN-DB and OSEAN-KG can be used to support different use cases.

### OSEAN-VIGET: The first SEA CDM use case for vaccine studies

As a first use case study of SEA CDM research, we focused on converting the VIGET^24^ data to the OSEAN database format. Specifically, the VIGET data includes 28 studies with 4,859 experimental samples. The information is stored in two files: one file contains gene expression data for the 4,859 samples, and the other contains metadata about these samples and studies.

**Figure 3** shows an example pattern for the vaccine-focused studies in the VIGET OSEAN DB with accompanying metadata entries that would be derived from each category. Vaccine studies typically consist of immune response experiments that differ based on which material is used for an intervention. The two most common differences between different experiments would be those that utilize an actual vaccine or a control (e.g., a saline solution). Following vaccination, a blood sample can be drawn from the targeted organism and is used to produce processed experimental samples for an assay. Each organism would be placed into a shared *Group*, which can be used to analyze the results of their respective assays. Some vaccine studies additionally monitor an organism for adverse events from the vaccination or their prognosis when exposed to disease.

**Figure 3.**
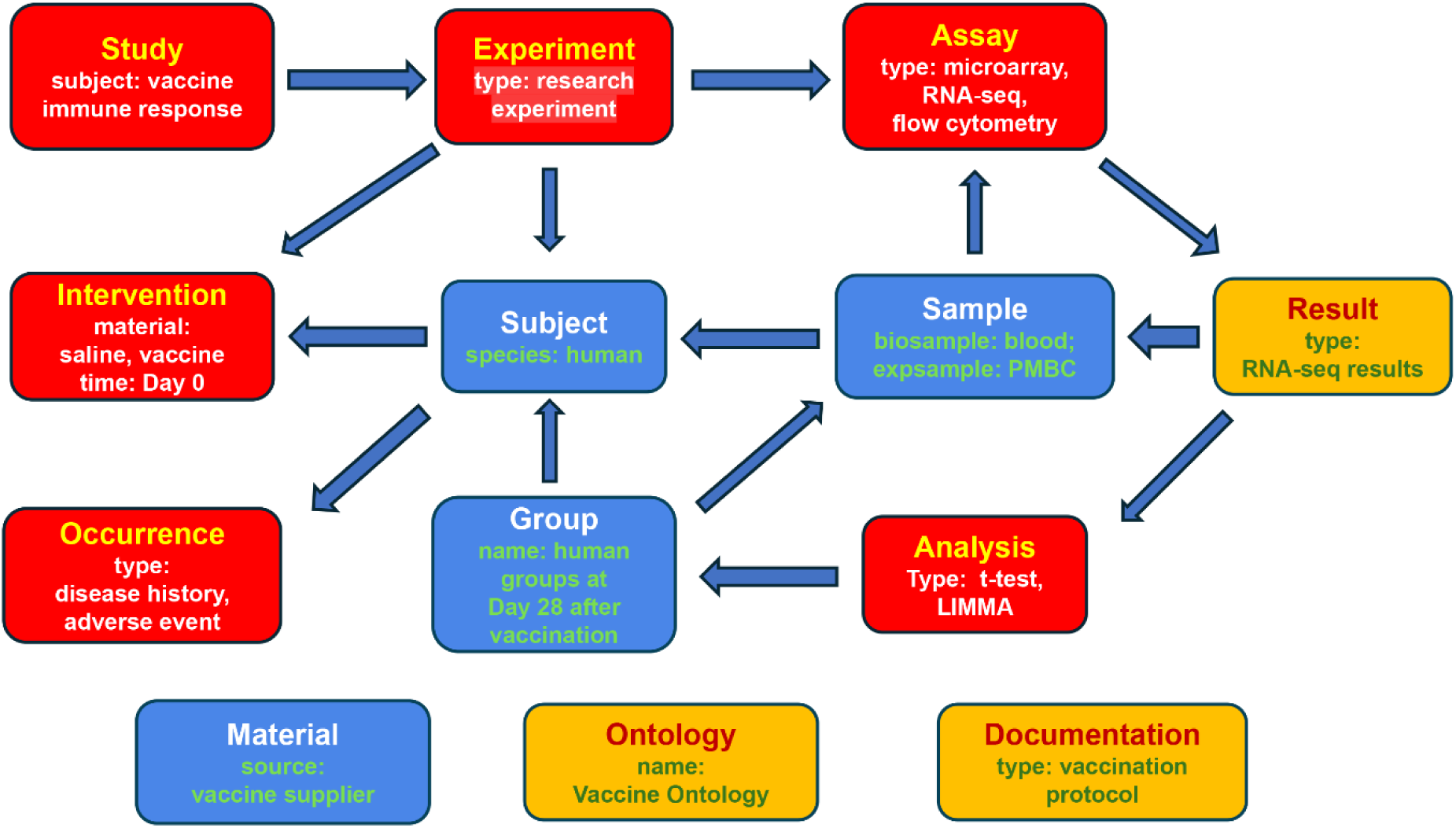
SEA CDM-based vaccine immune response study modeling. Vaccine studies exist within CELLxGENE, ImmPort, and VIGET. Vaccine studies can be focused on the organism’s immune response to a vaccine or can be focused on the organism’s response to disease following vaccination.

All the data in the two VIGET (metadata and gene expression) files were split, processed, and loaded into our VIGET OSEAN DB using an ETL (Extract, Transform, Load) pipeline, which is part of the PELAGIC Python module in the SEA CDM system. PELAGIC is a Python Engine for Linking, Analyzing, and Gleaning Insightful Context from the SEA CDM-based OSEAN databases. In addition to the ETL system, PELAGIC also includes other programs such as an OSEAN query and processing program as illustrated below.

### Querying VIGET OSEAN DB for scientific insights

The PELAGIC database query program includes the following features: querying the OSEAN database that contains standardized study metadata, setting up criteria for querying the results file(s), retrieving results from metadata and results file(s), and providing summarized results (**Supplemental Table 2**).

As a use case, the PELAGIC program was used to perform a simple query over the VIGET OSEAN database:

*Which genes are stimulated by the Fluarix flu vaccine in humans at Day 7 after vaccination?*

**Figure 4** provides the pipeline used to answer the above query using the OSEAN database. VIGET stores gene expression data for all samples found within it that are mapped to a list of sample reference names from GEO. Using PELAGIC as a Python wrapper, a SQL query was generated to retrieve a list of samples that match our criteria. Afterwards, additional gene restriction criteria were used to find a list of genes; we defined a gene as stimulated if it had at least a 2-fold expression in comparison to day 0. This query led to the identification of 35 genes (**Supplemental File 3**) stimulated by Fluarix in humans at Day 7 after vaccination.

**Figure 4.**
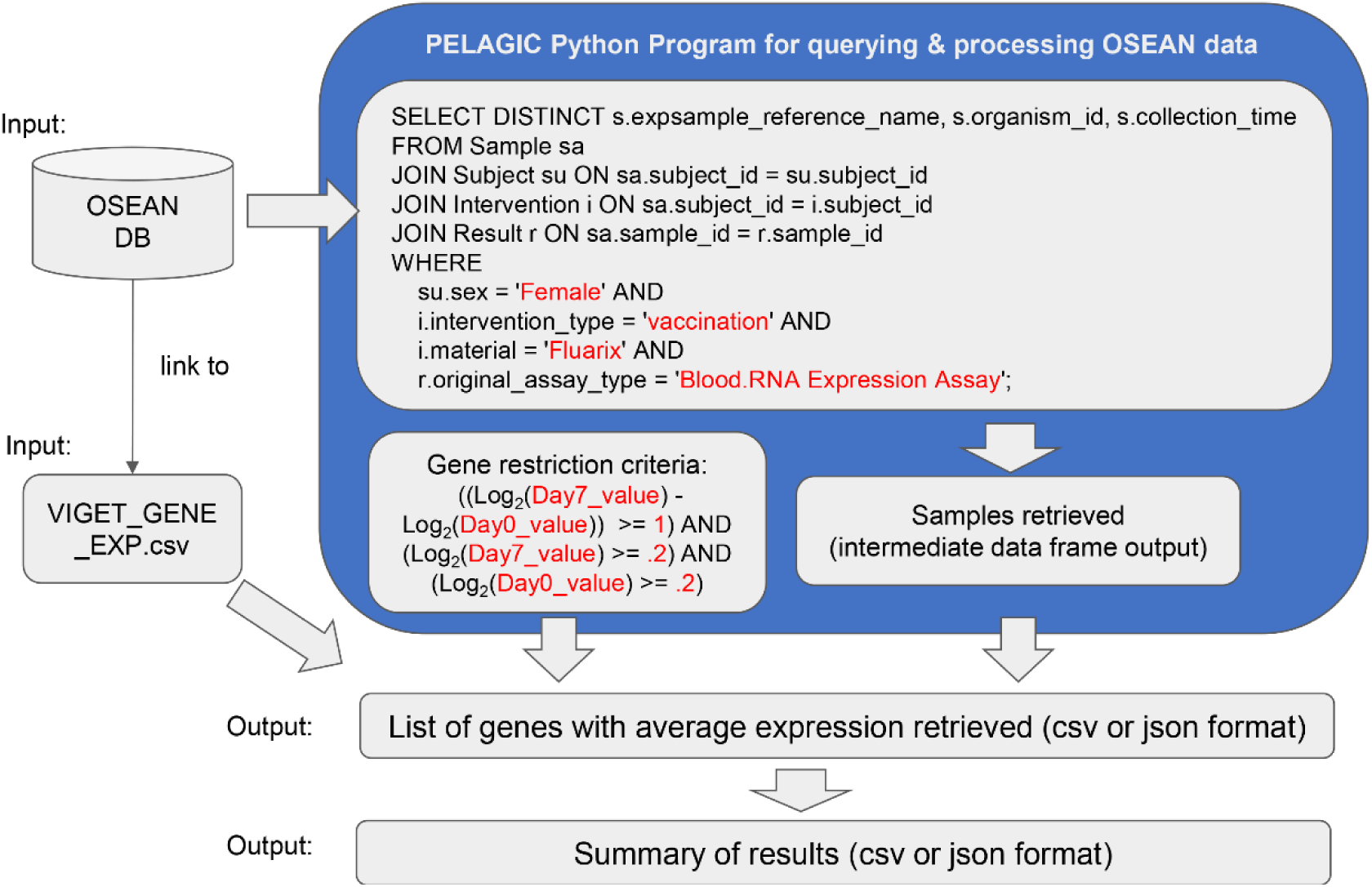
Example Query for VIGET OSEAN database. The blue box represents the PELAGIC Python wrapper used to query for data. Terms in red are variables that can be altered for querying different types of samples or genes. Three gene restriction criteria are included: at least 2-fold change (i.e., at least 1 in log2 value) of gene expression at Day 7 over Day 0, and gene expression value at Day 7 or Day 0 being at least 0.2 of log2. Furthermore, two additional criteria (at least 3 human subjects with the same gene profile and batch effect removed) were applied but not indicated in the figure (See more in Methods).

**Supplemental Figure 1** provides an alternative query that uses ontology IDs, which would result in the same outcomes as shown in **Figure 4**. In addition, the same data could also be represented and analyzed using SEA CDM-based knowledge graph instead of a relational database (i.e., OSEAN-DB) as described in a later section of this manuscript.

We then used the same method to analyze multiple influenza vaccines, both individually and all together. This resulted in 590 genes in total, with the breakdown of genes from the different influenza studies contained within VIGET **(Table 2)**. The gene overlapping results among individual influenza vaccines can be found as part of **Supplemental File 4**. The full lists of gene names, average, and values for the genes found in **Table 2** can be found in **Supplemental File 4.**

**Table 2.**
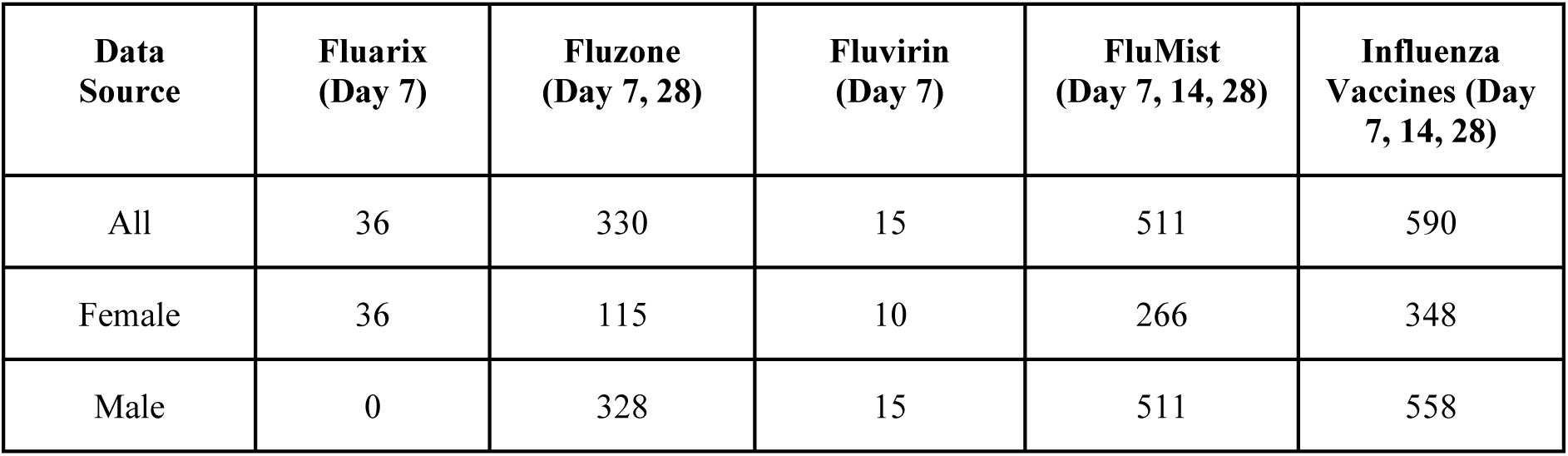
Influenza vaccine-induced gene response in humans as analyzed via OSEAN. The column of Influenza vaccines only covers the four listed vaccines as part of this table.

| Data Source | Fluarix (Day 7) | Fluzone (Day 7, 28) | Fluvirin (Day 7) | FluMist (Day 7, 14, 28) | Influenza Vaccines (Day 7, 14, 28) |
| --- | --- | --- | --- | --- | --- |
| All | 36 | 330 | 15 | 511 | 590 |
| Female | 36 | 115 | 10 | 266 | 348 |
| Male | 0 | 328 | 15 | 511 | 558 |

Following our collection of genes stimulated by influenza vaccines, we noted a reduced number of genes found in Fluarix and Fluvirin. To consolidate our gene lists, the four vaccines were grouped into an ‘inactivated (or killed) influenza vaccine’ (TIV) or ‘live attenuated influenza vaccine’ (LAIV) using VO. As such, the gene lists of the Fluarix, Fluzone, and Fluvirin vaccines were combined into the TIV gene list, while FluMist was the only LAIV.

With the above classification, we can further analyze the differential patterns of responses stimulated by LAIV and TIV in all human subjects, or female- or male-specific human subjects (**Table 2 and Figure 5A**). Overall, LAIV appears to stimulate more genes than TIV in all vaccinated human subjects or in vaccinated females or males, suggesting more active responses stimulated by LAIV compared to TIV. Furthermore, females appear to be associated with fewer influenza vaccine-stimulated genes compared to males.

**Figure 5.**
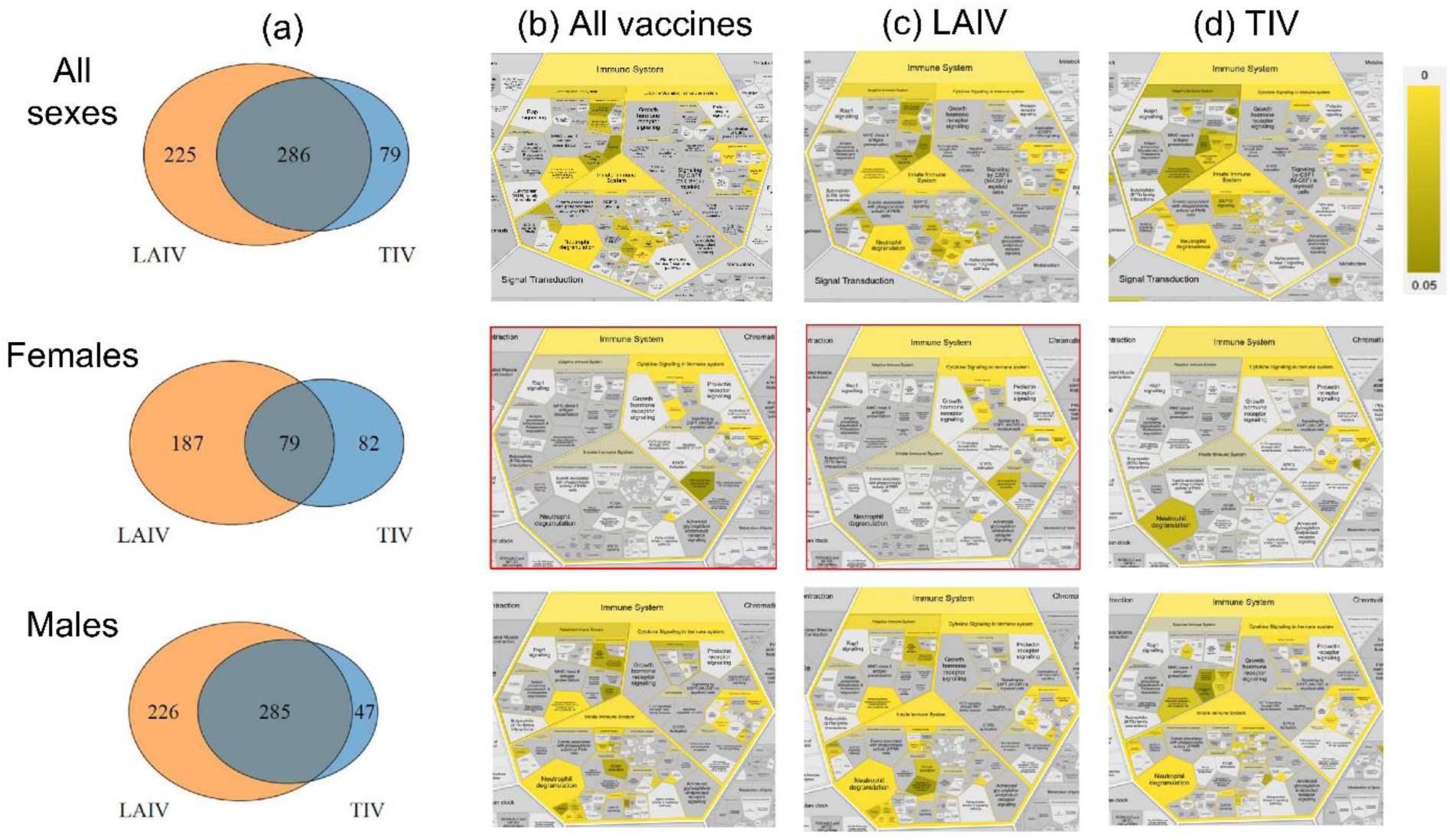
Statistically differential genes and associated immune response profiles in humans vaccinated with LAIV and TIV based on VIGET OSEAN data analysis. (a) Venn diagrams of the lists of differential genes in female, male, or all human subjects stimulated by LAIV and TIV. (b-d) show enriched Reactome immune system pathway profiles in female, male, or all human subjects stimulated by different vaccines, in which the combined LAIV/TIV-stimulated gene lists are used in (b), LAIV-stimulated gene lists are used in (c), and TIV-stimulated gene lists are used in (d). Boxes in red contain gene lists that did not have any statistically significant neutrophil degranulation or interleukin-4 and 13 signaling. The Reactome web tool ^25^ was used in the generation of (b-d) figures. **Supplemental File 4** contains the CSV files containing the full results of the Reactome functional analysis for this figure. Full version of the pathway images from (b-d) figures can be found in **Supplemental Figures 2-10.**

By examining the differential human gene lists stimulated by different influenza vaccines, we identified shared or vaccine-specific immune response pathways among all human subjects, females, or males (**Figure 5B-D**). For example, using all the combined genes found in human subjects of all sexes stimulated by all or individual influenza vaccines, we found many shared statistically significantly differential innate, adaptive, and cytokine signaling immune pathways, such as interleukin-10 signaling (False Discovery Rate (FDR) adjusted (the same below) p-value = 1E-5), ‘interleukin-4 and interleukin-13 signaling’ (p-value = 3.37E-7), and neutrophil degranulation (p-value = 1.1E-2). However, not all the pathways remained significant in sex-stratified sets. With all influenza vaccines combined, vaccinated males appeared to have similar immune profiles compared to the all-sex human subjects, and vaccinated females appeared to have less stimulated immune responses compared with males (**Figure 5B**).

Notably, LAIV-vaccinated females appeared to have unique immune responses compared to other vaccine groups (**Figure 5**). One common immune response relates to neutrophil degranulation, a process in which neutrophils release antimicrobial granules^26^. When all influenza vaccines and all sexes are considered, neutrophil degranulation was observed (**Figure 5B**), suggesting that influenza vaccines stimulate the release of antimicrobial granules from neutrophils. Such a phenomenon is also observed in males treated with all vaccines and females treated with TIV. However, neutrophil degranulation appeared to be missed in females vaccinated with live attenuated flu vaccine (e.g., LAIV). Meanwhile, we found ‘TNF binds their physiological receptors’ is uniquely enriched in LAIV-vaccinated females but not statistically enriched in LAIV-vaccinated males or in TIV-vaccinated female or male human subjects.

Neutrophil degranulation-associated differential genes in sex-specific human subjects were analyzed. A total of 33 and 13 genes were found to be differentially expressed in male and female human subjects, respectively, at Day 7 after LAIV vaccination (**Supplemental File 4**). Among them, six genes are shared: CEACAM8, CXCL1, FCAR, LAMP3, PLAUR, PSMD6, and PTX3. Examples of male-specific genes include SLC2A3, CD33, CD36, and LEC9. Examples of female-specific genes include CD47 and DDX3X.

In addition to the above Reactome pathway analysis, we also performed Gene Ontology (GO) enrichment^27, 28, 29^ (**Supplemental Figure 10**) and KEGG^30^ enrichment (**Supplemental Figure 11**). Both GO and KEGG analyses also showed more similarity in enriched functions between the all-sex and male gene sets than between the female gene set and either of the others. The female response to LAIV showed heightened patterns related to blood cells (‘erythrocyte development’ (FDR adjusted (the same below) p-value = 4.8E-7), ‘hemoglobin metabolic process’ (p-value = 4.8E-7). In contrast, TIV showed unique terms related to the down-regulation of genes related to stress response, such as ‘negative regulation of stress-activated MAPK’ (p-value = 1.03E-4). However, these sex differences tend to fade when the datasets are combined, with pathways related to inflammation becoming more prominent, including ‘regulation of immune response’ (p-value = 5.73E-10), ‘interleukin-1 production’ (p-value = 3.82E-9), and ‘myeloid leukocyte activation’ (p-value = 2.25E-9).

### SEA CDM-based OSEAN for storage and analysis of heterogeneous data from different resources

Following the conversion of VIGET into SEA CDM, we evaluated how the SEA CDM and its associated OSEAN database can be used to represent and store heterogeneous data from different resources. For this, we developed two use cases: one OSEAN database for storing the ImmPort data and another one for storing the CELLxGENE data.

Our ImmPort OSEAN database was able to convert all the ImmPort database contents to the SEA CDM format and store all the ImmPort metadata. The ImmPort database uses MySQL relational database format (ImmPort) and has 33 core tables and multiple linking tables that had to be consolidated into SEA CDM format. **Figure 6** provides a conversion strategy that converts 33 core tables to the SEA CDM format: for example, three intervention-related tables in ImmPort (i.e., “Intervention”, “Immune Exposure”, and “Treatment”) were merged into one Intervention table in SEA CDM. Likewise, three ImmPort sample-related tables (“Biosample”, “Control Sample”, “Expsample”) were merged into the Sample table in SEA CDM. A column called “type” in the Sample table is used to indicate these specific sample types. ImmPort has 15 tables to specify 15 specific assays. In SEA CDM, they were all merged into the Assay and Results tables in SEA CDM (**Figure 6**). **Supplemental Figure 12** and **Supplemental Table 3** include a clearer breakdown of the process used to convert the data. In the end, OSEAN-ImmPort stores all 1,263 studies (as of July 9th, 2025), which collected 1,650,290 experimental samples and 1,181,110 biosamples. All the ImmPort metadata was also loaded to the OSEAN-ImmPort database.

**Figure 6.**
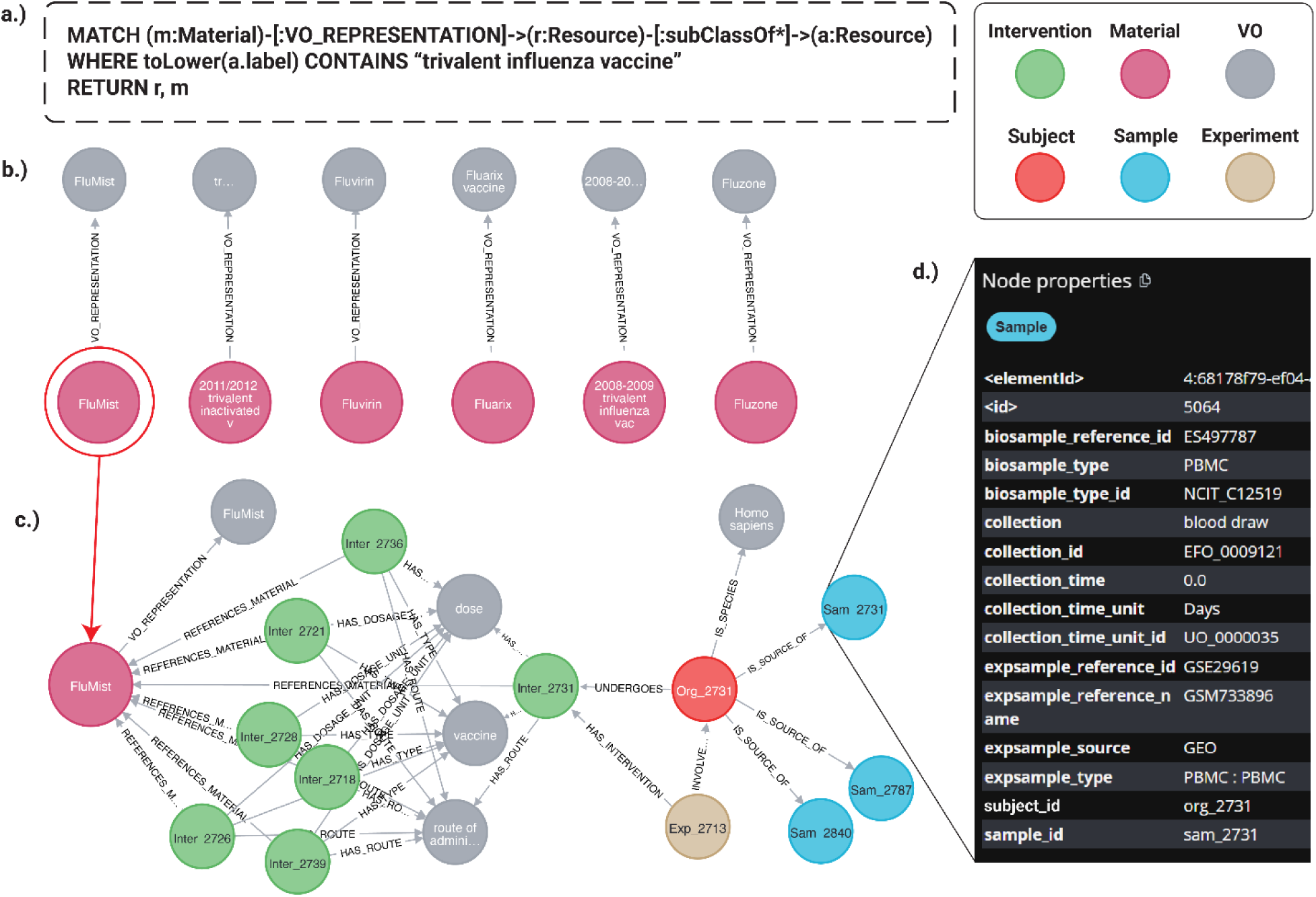
OSEAN KG representation of VIGET and VO attributes. A representative workflow for querying and navigating KG. **(a)** A Cypher query uses VO to find specific vaccines or materials related to “trivalent influenza vaccine.” **(b)** The results link the ontology term to specific entities like FluMist. **(c)** Expanding the FluMist node reveals its connections to experimental context, including Intervention and Sample nodes. **(d)** Selecting a sample node (Sam_2731) displays its detailed metadata, demonstrating the workflow from a broad query to a specific data point.

Our OSEAN database also successfully converted CELLxGENE data to SEA CDM format. CELLxGENE data are stored in the H5ad format^31^, a file format built on the HDF5 (Hierarchical Data Format 5) standard^32^, designed for storing large annotated scientific data. The H5ad format also stores variables and observations as different sections of the file. Specifically, CELLxGENE contains 10 common variables relating to organism, sample, occurrence, and results (organism, species, developmental_stage, disease, sex, tissue, cell type, suspension, assay, tissue_type) and 2 metadata (sample_id and observation_join_id) for each study **(Supplemental Table 4)**. Each common variable has its name and its ontology ID. A dedicated SEA CDM ETL was developed to convert the CELLxGENE columns to the SEA CDM format. In addition to these common variables, CELLxGENE also supports additional variables submitted by data providers, which requires specific attention.

In our demonstrative study, we focused on converting influenza-related CELLxGENE studies to OSEAN. Among the total of 1,844 CELLxGENE studies, we extracted 5 influenza-related datasets from 4 studies^33, 34, 35, 36^. Using the CELLxGENE website and the H5ad files downloaded from the website, 10 shared variables, along with 111 additional variables extracted from the H5ad files, were extracted from the five influenza-related datasets. Author-specific variables include additional intervention (tamoxifen treatment, O_2_ supplement), occurrences (comorbidities), or results from other assays (temperature, blood pressure). Note that initially separate OSEAN databases were developed for the ImmPort and CELLxGENE datasets. Since these OSEAN databases use the same SEA CDM schema, we found that these data can be easily merged into one single OSEAN database or use the same query program to query both databases.

With these OSEAN databases established, we asked one competency question: “*How many influenza-related studies and samples are in CELLxGENE and ImmPort?”* To address this question, we developed the following simple MySQL query, which can also run independently against the database or be part of the PELAGIC Python program:

SELECT COUNT(DISTINCT biosample_reference_name), COUNT(DISTINCT(study_id)), COUNT(DISTINCT expsample_reference_name)
FROM Sample sa
JOIN Subject su ON sa.subject_id = su.subject_id JOIN Intervention i ON i.subject_id = su.subject_id
JOIN Experiment e ON e.experiment_id = su.experiment_id
WHERE i.material IN (’influenza’, ’Influenza A virus’, ’Influenza A virus (A/California/7/2009(H1N1)’);

**Table 3** provides summarized results of this and other related queries. Specifically, ImmPort includes 228 influenza-related studies out of the total of 1,263 ImmPort studies on July 9th, 2025, representing over 18% of the ImmPort studies. Our study found 9,750 influenza-related expsamples and 48,567 influenza-related biosamples, with ImmPort containing a higher percentage of expsamples than compared to biosamples (4% and 1%, respectively). Our OSEAN query also found 42 biosamples and 149,469 influenza-related expsamples from the CELLxGENE OSEAN DB, which includes the five specific studies. Multiple biological and experimental sample types were identified (**Table 3**).

**Table 3.**
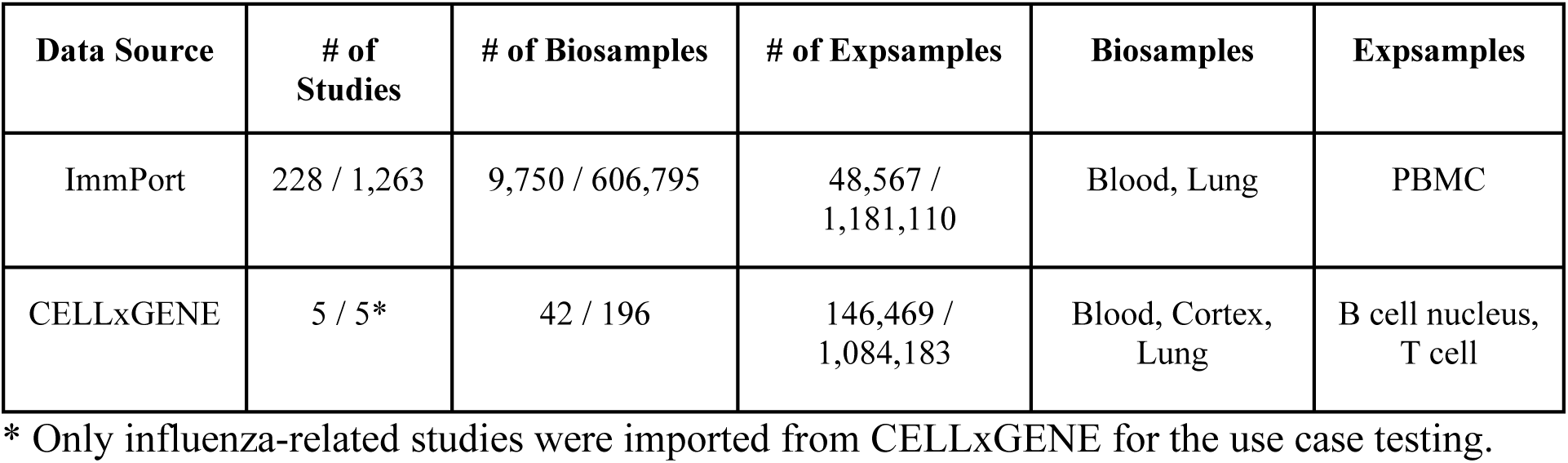
Influenza-related samples from two OSEAN DBs for ImmPort and CELLxGENE.

### SEA CDM-based knowledge graph creation and applications

We have generated a Neo4j knowledge graph to show how to integrate an ontology directly into the OSEAN DB related to VIGET. **Figure 6** illustrates a representative workflow within our Neo4j knowledge graph, demonstrating the powerful integration of ontological definitions with VIGET data. The process begins with a semantic query formulated in Cypher (**Figure 6A**), designed to identify all Material nodes associated with “trivalent influenza vaccine.” A critical component of this query is the use of a variable-length path, [:subClassOf*], which traverses the imported VO hierarchy. This feature enables a comprehensive search that retrieves not only direct subclasses but also all subclasses of the target term. The specific query shown in **Figure 6A** only includes nodes that overlap between the OSEAN VIGET DB and VO, dropping the restriction of matching the Material nodes would have returned over 700 different “trivalent influenza vaccine” nodes.

The initial results of this query are displayed in **Figure 6B**, returning a set of nodes that include specific and categorized products such as FluMist, bridging the gap between abstract ontological concepts and concrete entities within the database. **Figure 6C** showcases the exploratory capability of the graph. By expanding the FluMist node, the users can navigate its network of connections, revealing links to related entities such as the Intervention administered, the Organism studied (*Homo sapiens*), and the Experiment in which it was used.

This traversal continues further, as shown by the selection of the Sample node Sam_2731. The associated metadata for this specific sample is presented in **Figure 6D**, providing researchers with immediate access to critical details like its expsample_type (PBMC), collection method (blood draw), and the source GEO accession number (GSM733896). This seamless workflow, from a high-level conceptual query to detailed sample-level attributes, exemplifies how the knowledge graph empowers researchers to efficiently discover and interrogate complex, interconnected datasets.

## Discussion

In this manuscript, we present the “Study-Experiment-Assay” common data model (SEA CDM) together with the OSEAN database and PELAGIC program for seamless integration of heterogeneous data from biological and biomedical resources. We demonstrate the system across three data resources (i.e., VIGET, ImmPort, and CELLxGENE) and show that the simple SEA CDM system, supported with interoperable ontologies, can efficiently annotate, integrate, store, query, and analyze diverse datasets. Using our system, we successfully integrated and analyzed influenza vaccine-induced immune responses in human subjects, leading to the identification of various enriched immune profiles in females and males vaccinated with live attenuated and inactivated influenza vaccines.

SEA CDM is a simple but powerful solution founded on the common elements in biological and biomedical experimental studies. There are many standards that focus on specific domains, such as the MIAME standard for microarray experiments^37^, MIFlowCyt standard for defining the minimum information about a flow cytometry experiment^38^, MIRAGE standard for glycomics experiments^39^, and the ImmPort data model for immune response data standardization and storage. However, none of these standards or data models can be directly used to cover all the biomedical experimental studies. In contrast, SEA CDM captures the most basic elements from various biological and biomedical experimental studies. Every biomedical experimental study aims to answer a scientific question through one or more specific experiments. Each experiment typically generates samples that are further used by some lab assays. These three elements, *Study*, *Experiment*, and *Assay*, form the basis of the SEA model. Further exploring these three “process” level elements, we naturally identify *Organisms* and *Samples*, which are classes of “material entity” in an ontological sense^11^. We used “Organism” instead of “Person” (like in OMOP CDM) because our studies often include non-human organisms such as animals (e.g., mice) and bacteria. In addition, our SEA CDM includes *Group*, *Analysis*, and *Result* to cover the basic needs of biomedical studies for statistical analysis. In comparison, the OMOP CDM does not include *Group* and *Analysis*, as it focuses on individual patients rather than studies with grouped organisms. Although they appear simple, these elements form the fundamental parts of our SEA CDM system and can cover various types of studies.

While the basic structure of SEA CDM is simple, it is powerful in its usage of interoperable ontologies in data standardization and inference. The simple SEA CDM model may make it difficult to cover a lot of details about specific entities, such as vaccines. However, the solution to this limitation is using interoperable ontologies that store not only the standardized terms but also the knowledge behind these terms. For example, for a specific vaccine like “FluMist”, if we have its VO identifier, we can easily search and find that it is a live attenuated vaccine against influenza viral infection and that it is designed for humans. Ontologies have been used successfully to standardize investigations from the ISA framework^14^ and from electronic health records within OMOP CDM^16^. There are two different ways of using ontologies. One way is to directly include all the ontology terms in the system (like OMOP CDM or what our OSEAN-KG demonstrated); and the other way is to call another external ontology triplestore database or knowledge graph like Ontobee^40^, which is what we did in our OSEAN-DB demonstration (**Figure 2B**). Overall, the incorporation of interoperable ontologies in the SEA CDM system supports advanced integration and data FAIRness^9^. We may evaluate how we can use an OMOP-like method in ontology incorporation for data standardization in the future.

To our best knowledge, our proposed SEA CDM model is the first common data model tailored for standardizing and integrating various biological and biomedical experimental studies in a simple but solid relational database or knowledge graph format. The ISA framework^14^ and their associated ISA-TAB tools^15^ provide a flexible semantic framework and a set of web tools to support the standard representation of biomedical investigations. However, the ISA format lacks the explicit focus on experiments, and its semantic representation of various investigation types appears too general to be used. The ImmPort data model is well developed, with over 30 tables, to cover the specifics of immune response studies. However, the model is often too narrowly focused on immunological studies and does not fit the broad coverage of the SEA CDM. The OMOP CDM is broadly used for standardization and integration of electronic health records (EHR). It has many advanced features such as a simple schema and the usage of standardized vocabulary. While the OMOP CDM focuses on individual patients’ records, it is not suitable for standardization and integration of various biomedical experimental studies.

The SEA CDM has been applied successfully in both relational database and knowledge graph formats. We started our testing with the relational database format by following the usage of the OMOP CDM and ImmPort data model. The practice in the OMOP CDM database and ImmPort database has been proven successful. Our own practice in the SEA CDM-based OSEAN databases also shows its solid compatibility. The reasons behind these successes are likely due to their simple, structured design and easy-to-query ability. Meanwhile, we tested the usage of SEA CDM in knowledge graph generation. Our results show that it provides additional benefits, such as the natural, easy inclusion of VO and the easy visual exploration of the graph classes and attributes. We think both formats can be used for different needs. While they can be complementary, each format can be independently implemented and does not need to rely on the other.

Using our SEA CDM and its associated OSEAN-DB system, we successfully integrated and analyzed influenza vaccine-induced immune responses in human subjects, leading to the identification of distinct immune profiles in females and males vaccinated with various influenza vaccines. For example, our study found while neutrophil degranulation was observed in males treated with all vaccines and females treated with inactivated influenza vaccines, the phenomenon was not found in females vaccinated with live attenuated flu vaccine (e.g., LAIV). As the most abundant human white blood cell, neutrophils provide a first line of innate immune defense. Through neutrophil degranulation, neutrophils release the contents of their intracellular granules (enzymes, antimicrobial peptides, and reactive oxygen species) into the extracellular space or phagosome in order to kill pathogens. Our male-biased finding aligns with the recent discovery in a mouse model that male neutrophils tend to exhibit higher degranulation activity than females^26, 41^. To our best knowledge, our results represent the first report of sex-biased while neutrophil degranulation in responses to live attenuated influenza vaccines, which may contribute to the sex-biased responses to LAIV^42^. Our previous research also found different adverse event profiles in human responses to LAIV and trivalent inactivated influenza vaccines (TIV) as reported earlier^43^, some of which may also be explained by the neutrophil degranulation and other differences found in our research. Therefore, our findings support better understanding of the sex effect on the influenza vaccine immunology and supports rational vaccine design and dosing strategies.

While a biological question is often complicated, our SEA CDM system provides an effective way to tackle nuanced contexts such as temporal dynamics, comorbidities, and drug interactions. Our reported use case lays out how different dates after vaccination could be used in our queries to support the analysis of temporal dynamics of host immune responses to vaccines. Meanwhile, SEA CDM provides mechanisms to study other contexts. SEA CDM includes the class Intervention, which can be used to study the effects of specific interventions such as drugs and vaccines. Potential drug-drug interactions can be studied by looking for Interventions that contain multiple drug administrations to the same organism that occurred at the same time or if the second drug administration is done before the first drug has been metabolized. The Occurrence class in SEA CDM can further be used to support the study of comorbidities. For example, the ImmPort entries also contain information related to the presence of specific comorbidities. These results can be queried by examining the Occurrence class.

Our future work will proceed in several directions. First, different elements of the SEA CDM can be further standardized. For example, we plan to standardize each assay, experiment, and even study with standardized protocols. The protocols.io framework^44^ provides an integrative platform to standardize protocols with their associated reagent materials, assay platforms, parameters, or methods (i.e. formulas). Our SEA CDM can refer to these protocols in the protocols.io to support reproducible experimental studies. Like ontology providing implicit knowledge, the reference of these protocols provides foundational knowledge for individual studies. We can also develop SEA CDM-based OSEAN databases for integrating data from other resources, such as TCGA^6^, KPMP^7^, and HubMAP^8^. In addition, to better support ontological standardization and integration of the SEA CDM metadata and experimental attributes, we are developing an ontological representation of the SEA CDM metadata using the community-based Ontology of Precision Medical Intervention (OPMI) as the platform ^45, 46, 47^.

Furthermore, it is possible to integrate all these datasets under the SEA CDM framework, so that we can systematically analyze all these heterogeneous datasets simultaneously. To make the whole integrative system work properly, we will also further enhance the PELAGIC module into a tool suite to include database-specific ETLs to convert database-specific columns into their SEA-CDM equivalents. Command-line tools and help options will be developed to aid in mapping author-labeled metadata into SEA-CDM. One module will include checking duplicate entries for specific rows that are shared between databases. Such functionality could include an option to add programs that call upon protocol.io and tag the entries using the Bioschemas format^48^. There is no comprehensive list of data formats or protocols that can be linked to OSEAN DB. Ultimately, expanding OSEAN to store these findings will demonstrate that all models can flow into SEA CDM.

## Methods

### General SEA CDM modeling method

A hybrid approach combining both top-down and bottom-up methodologies was utilized to develop the SEA CDM Model. For the top-down modeling, we investigated different models, including ISA format^14^, ImmPort Data Model^1^, OMOP^16^, and VIO modeling^49^, and incorporated or mapped many features of these models to SEA CDM. For the bottom-up modeling, we analyzed specific vaccine or influenza studies as reported in VIOLIN, ImmPort, and CELLxGENE to guide our SEA development. During the SEA CDM development, we identified a minimal set of metadata types that are required to fully present individual studies. The basic rule of thumb is that without such metadata, the report of the biological experiment is incomplete, and others will not be able to repeat the study.

### Ontological representation of SEA metadata types and their relations

After SEA metadata types were identified, we mapped these metadata type terms to ontology terms. As our VIGET use case focused on host immune response to vaccines, the Vaccine Ontology (VO)^50^ was used as the default ontology and platform for the ontology term mapping. For other ontologies, we used Ontobee^40^ to identify the original ontology term ID to help guide the assignment of different column entries. The following additional ontologies were used to help define appropriate ontology IDs for different metadata categories: Cell Ontology^51^, Disease Ontology^52^, Experimental Factor Ontology^53^, Gene Ontology (GO)^28, 29^, Human Phenotype Ontology^54^, NCBI Taxonomy^55^, NCI Thesaurus^56^, Protein Ontology^57^, Ontology of Biomedical and Clinical Studies^58^, Ontology of Adverse Events^59^, Ontology of Biomedical Investigations^13^, Uberon Anatomy^60^, and Unit Ontology^61^. A high-level ontological hierarchy was also designed to cover all the identified metadata types based on the VO overall structure. The Protégé OWL editor^62^ was used for ontology visualization.

### Development of SEA CDM-based OSEAN relational databases

The SEA CDM-based database schema was used to generate three specific MySQL databases, which are all named as OSEAN (Ontology-based Study-Experiment-Assay Network) databases (DBs). MySQL Workbench 8.0 was used for database management and queries. These three OSEAN DBs were populated using the data extracted from existing resources. Customized automatic Extract-Transform-Load (ETL) pipelines were developed to expedite the OSEAN DB generation.

The first OSEAN DB developed was called OSEAN-VIGET, which converted the data from the VIGET^3^ (Vaccine Immune Gene Expression Tool) program to the SEA CDM format. Specifically, two VIGET input files were extracted from the VIGET Zenodo website (https://zenodo.org/records/7407195), with one about the metadata of 28 vaccine immune studies, and the other containing the normalized gene expression data file. A Python ETL program was developed to process the metadata file into the SEA CDM format. The gene expression data file was downloaded to a local directory pointed to by the SEA CDM documentation.

A designated Zenodo repository (https://zenodo.org/records/17770032) was also created to store SEA CDM and OSEAN-DB related resources, including SQL code for generating the OSEAN MySQL, SEA CDM-based spreadsheet templates for populating the database, and Python scripts for transforming VIGET files into OSEAN-DB format. The VIGET example represents our first complete demonstration for the SEA CDM and OSEAN-DB workflow. The materials provided in the Zenodo repository enable full reproduction of the work.

The second OSEAN DB is OSEAN-ImmPort, which collected all metadata from the ImmPort^1^ database by downloading the information from the ImmPort web server (version used: ALL_STUDIES_DR 56.2)^1^.

OSEAN-CELLxGENE is the third OSEAN DB, which stores the metadata of five influenza datasets from four studies^33, 34, 35, 36^ deposited in the CELLxGENE database (<u>cellxgene.czisience.com</u>)^5^. The data from these four studies^33, 34, 35, 36^ are stored in five separate H5ad files, which were identified after searching for ‘influenza’ in CELLxGENE. A Python ETL program was developed to parse these H5ad files and store the data in the SEA CDM format.

### PELAGIC Python module developed for OSEAN data query and analysis

Our PELAGIC module is a Python engine program for linking, analyzing, and gleaning insightful context from the SEA CDM-based OSEAN databases. PELAGIC was developed for querying and analyzing the data from OSEAN databases. PELAGIC was coded in Python version 3.12. The first module contains a list of functions used by modules, including the creation of SEA-CDM templates for different SEA classes and foreign keys for OSEAN DB into dataframes. The second set of modules is generic ETL modules used to convert CELLxGENE, ImmPort, and VIGET. The third set of modules related to querying the OSEAN database allows the user to input customized queries into OSEAN-DB to export the results as a CSV or JSON file.

### OSEAN-based analysis of Influenza vaccine-induced gene expression profiles

We explored the metadata and gene expression data of VIGET^3^ using a custom Python script as part of the PELAGIC module. First, a specific MySQL query embedded in the Python script was developed to query the OSEAN database for a set of samples that met specified query criteria. Second, the Python program also includes a set of customizable gene expression restriction criteria for filtering samples based on the locally stored gene expression data file. In addition, to enhance the robustness of the data analysis, any genes that had less than three samples after the above filtering were rejected, and the samples listed as having a batch factor within the VIGET data file were also removed from the gene set expression analysis.

The gene lists were compiled individually for Fluzone, Fluarix, FluMist, and Fluvirin. The FluMist and Fluzone selection included vaccines that were labeled as ‘live attenuated vaccine’ and ‘trivalent influenza vaccine,’ following how VIGET classifies these vaccine categories. Venn diagrams were generated to visualize the overlaps and unique gene subsets across vaccine groups and genders. GO and Kyoto Encyclopedia of Genes and Genomes (KEGG) pathway enrichment analyses were performed using our in-house R package richR (https://github.com/hurlab/richR) to identify significantly enriched biological processes and pathways within each subset of the Venn diagrams. A custom R script was developed to automate the workflow and generate summary plots and tables for downstream interpretation. The resulting gene lists were then used on the Reactome website^25^ for further functional and pathway analysis.

### Development of SEA CDM-based knowledge graph using Neo4j

The entire database, along with the Vaccine Ontology (VO, version 2025-07-06), was imported into Neo4j using the Neosemantics (n10s) plugin^63^ and a custom Python script (https://github.com/sea-cdm/OSEAN-KG/blob/main/Import_to_neo4j.py). The setup began with configuring Neosemantics (n10s) to support effective ontology integration by defining rules for handling Uniform Resource Identifiers (URIs), relationship types, and RDF type semantics. The VO ontology file was then loaded, which was automatically parsed into RDF triples. These triples were stored as Resource nodes, establishing the core vocabulary of VO within the knowledge graph. Following the import, mapping procedures were applied to connect VIGET and ImmPort nodes to VO entities, enriching the ontology with relevant relationships and properties from the integrated datasets.

### Code Availability

All code related to the SEA CDM program is available at https://github.com/SEA-CDM. Three repositories are deployed, including the general SEA CDM repository that provides the overall SEA CDM design and documentation, the OSEAN-DB repository that provides the OSEAN MySQL relational database system, and the OSEAN-KG repository for the OSEAN knowledge graph system using Neo4j.

The OSEAN-DB repository (https://github.com/SEA-CDM/OSEAN-DB) includes the code of the MySQL schema for loading OSEAN DB, the ETL programs for extracting, transforming, and loading data from different resources to OSEAN DB, the functions of the PELAGIC Python module, and the three use case OSEA DB studies.

The OSEAN-KG repository (https://github.com/SEA-CDM/OSEAN-KG) includes the code for generating a VIGET Neo4j knowledge graph using SEA CDM, the .csv data files containing non-empty tables generated via the VIGET ETL, and Python functions for importing data to Neo4j and ontology mapping.

The static web application, accessible at https://sea-cdm.github.io/SEA-CDM/, was deployed via GitHub Pages from the repository’s main branch (https://github.com/sea-cdm/SEA-CDM). The GitHub repository used an organized directory structure for resources, documentation, and core files. A custom template.js script detected the base path, corrected links, and injected shared interface components to ensure consistent navigation across local and hosted environments. This approach enabled maintainable and modular development.

### Data Availability

The SEA CDM GitHub organization website (https://sea-cdm.github.io/SEA-CDM/) provides details and code related the standards of SEA CDM (https://github.com/sea-cdm/SEA-CDM), OSEAN-DB (https://github.com/sea-cdm/OSEAN-DB), and OSEAN-KG (https://github.com/sea-cdm/OSEAN-KG). All data used for the three OSEAN DBs (OSEAN-VIGET, OSEAN-ImmPort, OSEAN-CellxGene) and OSEAN-VIGET-KG are extracted from the openly available data sources VIGET^3^, ImmPort^1^, and CELLxGENE^5^. The exact methods for the extraction and usage of these datasets are provided in the README file for the OSEAN-DB GitHub repository (https://github.com/sea-cdm/OSEAN-DB/blob/main/README.md). The Zenodo repository (https://zenodo.org/records/17770032) stores more data related to this manuscript.

## Acknowledgements

We appreciate the discussions about the SEA CDM system when it was presented as a poster in the Data Repositories and KnowledgeBases (DRKB) 2025 Program Network Meeting and as an oral presentation at the Intelligent Systems for Molecular Biology (ISMB) 2025 Conference. We had some minor use of ChatGPT during discussions, as well as during classifications of vaccine studies and creation of the PELAGIC module.

## Author Contributions

AH developed the OSEAN MySQL database and PELAGIC tools and implemented the use cases. FYY was responsible for OSEAN knowledge graph generation and establishing the GitHub pages. JH was responsible for GO and KEGG enrichment analysis. AH, JH, JZ, AMM, GW, CT, BA, and YH participated in SEA CDM design. YH initiated the overall project design and served as a vaccine domain expert. AH and YH took primary creation of figures and tables and prepared the first complete paper draft. All authors provided discussion, reviewed the manuscript, and approved the paper submission.

## Supplemental Materials

**Supplemental Table 1. General metadata tables and columns in OSEAN DB.** Each column indicates a variable or foreign ID. All variables are represented via a name and matched ontology ID. All metadata columns after the semi-colon are foreign IDs used by OSEAN DB.

**Supplemental Table 2. Conversion of VIGET metadata into OSEAN format.** Each box represents a table containing data related to VIGET. Class entries are listed by class.row format. Related SEA-CDM class entries represent additional annotations done to VIGET. One row, expsample_to_multiple_gsm_flag can be answered via querying the data in OSEAN and thus dropped.

**Supplemental Table 3. Mapping of common ImmPort core tables to SEA-CDM classes.**

**Supplemental Table 4. Mapping of common CELLxGENE metadata variables category to SEA CDM categories.**

**Supplemental Figure 1. Alternative Ontology-based Query for VIGET OSEAN database.** This figure provides an alternative query using the ontology IDs instead of the main queries as part of **Figure 4**. These queries are identical due to prior data harmonization ensuring that each attribute pair (name, ontology_id) is a 1:1 match. The time units were not included as part of **Figure 4** for legibility. The mapping for each figure and ontology ID is as follows: ‘PATO_0000383’ is ‘female’; ‘VO_0000002’ is ‘vaccination’; ‘VO_0021053’ is FluMist; ‘OBI_0001985’ is ‘microarray assay’; and ‘UO_0000033’ is ‘day’.

**Supplemental Figure 2. Full Reactome representation of Reactome pathways stimulated by all Influenza vaccines in all-sex human subjects.** Gene set enrichment values can be found as part of **Supplemental File 4**.

**Supplemental Figure 3. Full Reactome representation of Reactome pathways stimulated by all Influenza vaccines in female human subjects.** Gene set enrichment values can be found as part of **Supplemental File 4.**

**Supplemental Figure 4. Full Reactome representation of Reactome pathways stimulated by all Influenza vaccines in male human subjects.** Gene set enrichment values can be found as part of **Supplemental File 4.**

**Supplemental Figure 5. Full Reactome representation of Reactome pathways stimulated by live attenuated Influenza vaccines in all sexes.** Gene set enrichment values can be found as part of **Supplemental File 4.**

**Supplemental Figure 6. Full Reactome representation of Reactome pathways stimulated by all Influenza and live attenuated Influenza vaccines in female human subjects.** Gene set enrichment values can be found as part of **Supplemental File 4**.

**Supplemental Figure 7. Full Reactome representation of Reactome pathways stimulated by live attenuated Influenza vaccines in male human subjects.** Gene set enrichment values can be found as part of **Supplemental File 4**.

**Supplemental Figure 8. Full Reactome representation of Reactome pathways stimulated by trivalent inactivated Influenza vaccines in all-sexes.** Gene set enrichment values can be found as part of **Supplemental File 4**.

**Supplemental Figure 9. Full Reactome representation of Reactome pathways stimulated by trivalent inactivated Influenza vaccines in female human subjects.** Gene set enrichment values can be found as part of **Supplemental File 4**.

**Supplemental Figure 10. Full Reactome representation of Reactome pathways stimulated by trivalent inactivated Influenza vaccines in male human subjects.** Gene set enrichment values can be found as part of **Supplemental File 4**.

**Supplemental Figure 11. GO functional analysis results of influenza vaccines.** Any pathway listed shows up as one of the top 10 most significant GO pathways for one of the nine gene sets.

**Supplemental Figure 12. KEGG functional analysis results of influenza vaccines.** Any pathway listed shows up as one of the top 10 most significant KEGG pathways for one of the nine gene sets.

**Supplemental Figure 13. ImmPort to SEA-CDM format modeling.** A simplified mapping of key tables of ImmPort to SEA-CDM format. SEA-CDM foreign IDs require information that is found in linking tables to consolidate data (i.e., SEA-CDM Sample requires data loaded from the “Biosample”, “ControlSample”, “ExpSample” core tables and “Biosample-2-Expsample”. Additionally, information related to the SEA-CDM Sample’s Organism would require the use of the “Biosample-2-Subject” table. The original connections showing each table were taken from the ImmPort website.

**Supplemental File 1**. **Master SEA CDM Template File**. This file contains pre-generated SEA CDM template files and documentation for each SEA CDM class attribute. Bolded entries represent the default Ontology ID for an attribute that all Ontology IDs must have as an ancestor.

**Supplemental File 2. OSEAN-DB Instruction File**. This instruction file describes how to access to all data and code to generate OSEAN-DBs or OSEAN-KGs from existing SEA CDM files. We have also included the template files for OSEAN VIGET DB, OSEAN ImmPort DB, OSEAN CellxGene DB, and OSEAN VIGET KG.

**Supplemental File 3**. Summary of gene expression data from VIGET PELAGIC query.

**Supplemental File 4**. Summary of gene set expression analysis data and functional analysis from VIGET PELAGIC query.

