## Supplemental File 1 for "SEA CDM: Study-Experiment-Assay Common Data Model and Databases for Cross-Domain Data Integration and Analysis"

**Supplemental File 1. CDM model template instruction**

This Supplemental File includes links to the csv files and example usage of SEA-CDM terms for SEA CDM version 1.3.

**Study**

Data input Template:

<https://github.com/sea-cdm/OSEAN-DB/blob/main/DB-Schema/study.csv>

Instruction:

<https://sea-cdm.github.io/SEA-CDM/sea-cdm_documentation/study.html>

**Supplemental Table 1**.

| **Property** | **Description** | **Example Usage** |
| --- | --- | --- |
| study_id | A primary key used to represent a study. | “STUDY_1” |
| study_name | A string representing the name of a study. | “OSEAN-DB Vaccine Gene Response Study” |
| study_description | A string representing a short description of a study. This is set to 500 characters or less and is intended to illustrate the key question of a study. | “A study to analyze differences in vaccine gene response data from influenza using data in OSEAN” |
| study_type | A string representing the type of a study. | ‘Clinical Investigation’, ‘Experimental Study’ |
| study_type_id | An ontology string representing the type of a study. | OBI_0003697 (clinical investigation),  OBI_0500000 (study design) |
| study_focus | A string representing the intended process a study is about. | ‘Immune Response’, ‘Vaccine Response’ |
| study_focus_id | An ontology string representing the study focus. | Immune response (GO_0006955),  Vaccine response (VO_0005433) |
| Study_keywords | A string representing the keywords (usually diseases or qualities) to be used in the study. | Influenza, Tuberculosis |
| Study_keyword_ids | An ontology string representing the study keyword. | NCBITaxon_11320 (influenza), MONDO_0018076 (tuberculosis) |
| reference_source_id | A foreign ID used to represent the source of a study from another source. Must be paired with reference_source. | ‘SDY562’ |
| reference_source | A string representing the reference source for a study. | ‘ImmPort’ |
| comments | A string used as a placeholder for extra words or comments. |  |

SEA CDM Table:

**Experiment**

Data input Template:

<https://github.com/sea-cdm/OSEAN-DB/blob/main/DB-Schema/experiment.csv>

Instruction:

<https://sea-cdm.github.io/SEA-CDM/sea-cdm_documentation/experiment.html>

| **Property** | **Description** | **Example Usage** |
| --- | --- | --- |
| experiment_id | An primary key that corresponds to an instance of an experiment. | “EXPER_1” |
| study_id | A foreign key that corresponds to a study an experiment belongs to. | “STUDY_1” |
| documenation_id | A foreign key that corresponds to a document that describes the protocol used as part of this experiment. | “DOCU_1” |
| experiment_control | A binary flag used to indicate if an experiment row is used as part of an experimental control or not. | “True”, “False” |
| experiment_type | A string used to describe the type of an experiment performed. | “Clinical Visit”, “Experiment” |
| experiment_type_id | A string used to match experiment_type to an ontology ID. | “OBI_0500001” (clinical study design), “OBI_0500000” (study design) |
| experiment_subject | A string used to represent the type of subject used for an experiment. | Human, cell line |
| experiment_subject_id | A string used to match experiment_subject to an ontology_id. | “NCBI_Taxon9606” (human),  “CLO_0000031” (cell line) |
| reference_source_id | A foreign ID used to represent the source of an experiment from another source. Must be paired with reference_source. | “ARM_0000002” |
| reference_source | A string representing the reference source for an experiment. | ImmPort |
| comments | A string used as a placeholder for extra words. |  |

SEA CDM Table:

**Assay**

Data input Template:

<https://github.com/sea-cdm/OSEAN-DB/blob/main/DB-Schema/assay.csv>

Instruction:

<https://sea-cdm.github.io/SEA-CDM/sea-cdm_documentation/assay.html>

| **Property** | **Description** | **Example Usage** |
| --- | --- | --- |
| assay_id | An internal id used to describe an assay type. | “ASSAY_1” |
| documentation_id | An id of the documentation table used to identify the reference document describing the process of an assay. | “DOCU_1” |
| assay_name | A string representing the name of an assay; this specifies a general type of features. | “RNA-Microarray” |
| assay_name_id | A string representing the OBI ID for an assay. | “ECO_0000097” (RNA microarray) |
| assay_type | An assay category used to assign the type of the assay it belongs to. This is used to describe measurement or observation or survey. | “Experimental Assay”, “Observation” |
| organism_input | A binary value representing if an assay necessarily involves an organism as a specimen for an assay, instead of just using a sample. | True, False |
| reagents | A list of material IDs that are used as an input by an assay that are consumed by the process. This is used for experimental assays. | “Anti-mouse antibodies” |
| platform | A list of material IDs that are used as an input by an assay that are not consumed by the process. This is used for experimental assays. | “GPL5263” |

SEA CDM Table:

**Analysis**

Data input Template:

<https://github.com/sea-cdm/OSEAN-DB/blob/main/DB-Schema/analysis.csv>

Instruction:

<https://sea-cdm.github.io/SEA-CDM/sea-cdm_documentation/analysis.html>

| **Property** | **Description** | **Example Usage** |
| --- | --- | --- |
| analysis_id | A primary key used to identify a specific analysis. | “ANLYS_1” |
| group_id | A foreign key used to identify a specific group used for an analysis. | “GROUP_1” |
| documentation_id | A foreign key used to provide the documentation_id in the Documentation table that provides the analysis protocol and other related information used to describe an analysis. | “DOCU_1” |
| input_data | A string listed the data types that are used as inputs for a statistical analysis. | “VIGET” |
| input_data_id | A string listing the unique identifier for the input_data. | “https://viget.violinet.org/” |
| file_access | A string listing the download link / URL needed to access the file for the input data. | “https://zenodo.org/records/7407195” |
| analysis_name | A string representing the name of a type of analysis. | “LIMMA” |
| analysis_name_id | A string used to provide an ontology identification of an analysis name id. | “OBCS_0000168” |
| reference_source_id | A string used to identify an external reference for the group. | “37180100” |
| reference_source | A string used to identify the source of the reference_source_id. | “PMID” |

—------

SEA CDM Table:

**Intervention**

Data input Template:

<https://github.com/sea-cdm/OSEAN-DB/blob/main/DB-Schema/intervention.csv>

Instruction:

<https://sea-cdm.github.io/SEA-CDM/sea-cdm_documentation/intervention.html>

| **Property** | **Description** | **Example Usage** |
| --- | --- | --- |
| intervention_id | A primary key used to identify an experiment intervention. | “INTER_1” |
| experiment_id | A foreign key that is used to link an experimental intervention to an Experiment. | “EXPER_1” |
| subject_id | A foreign key that is used to link an experimental intervention to a Subject. | “ORGO_1” |
| material | A string used to identify the material used as part of an intervention. | “Afluria”, “Tamoxifen” |
| material_name_id | A string used to identify the ontology id that is associated with the material name. | “VO_0000006” (Afluria),  “CHEBI_41774” (Tamoxifen) |
| dosage | A string used to identify the dosage used of a material given to an experimental subject. | 0.25 |
| dosage_unit | A double used to identify the unit of a dosage. | “mL” |
| dosage_unit_id | An integer used to identify the unitID mapped to the dosage. | “UO_0000098” (milliliter) |
| intervention_type | A string used to identify the type of intervention. | “vaccination” |
| intervention_type_id | A string used to identify the ontology ID for route. | “VO_0000002” |
| intervention_route | A string used to identify the route of administration for an experiment intervention. | “Intravenous” |
| intervention_route_id | A string used to identify the ontology ID for intervention_route. | “VO_0000574” (intravenous) |
| t0_definition | A string used to identify when T0 is given for intervention. | “Time of administration” |
| intervention_time | A double used to identify how long after the start of T0 an intervention occurred. | 7.0 |
| time_unit | A string used to identify the unit for intervention_time. | “Day” |
| time_unit_id | A string used to identify the ontology ID for time_unit. | “UO_0000036” |
| reference_source_id | A string used to identify an external reference for the group. | “IMM00001” |
| reference_source | A string used to identify the source of the reference_source_id. | “ImmPort” |
| comments | Placeholder for additional comments. |  |

SEA CDM Table:

**Occurence**

Data input Template:

<https://github.com/sea-cdm/OSEAN-DB/blob/main/DB-Schema/occurrence.csv>

Instruction:

<https://sea-cdm.github.io/SEA-CDM/sea-cdm_documentation/occurrence.html>

| **Property** | **Description** | **Example Usage** |
| --- | --- | --- |
| occurrence_id | An internal identifier used to identify a specific occurrence. | **“**OCCUR_1” |
| subject_id | An identifier that maps to an experimental subject. | “ORGO_1” |
| occurrence_name | A string that lists the reported summary of an occurrence. | “Fever”, “Influenza” |
| occurrence_name_id | A string that lists the ontology ID used to identify an occurrence. | “HP_0001945”, “DOID_8469” |
| occurrence_severity | An integer that corresponds to the severity of an occurrence similar to the scale used by the Common Terminology for Adverse Events (CTCAE). Benign occurrences are rated as 0. | “0”, “1”, “2”, “3”, “4”, “5” |
| occurence_onset_date | A string used to identify the reference date for onset study. | “07-01-2025” |
| occurence_onset_datetime | A string used to identify the reference datetime for onset study. | “07-01-2025 14:02:56” |
| occurence_end_date | A double that lists the end of an occurrence for experimental subject. | “11-01-2025” |
| occurence_end_datetime | A datetime for the end of an occurrence. Leave this black if an occurrence is ongoing. | “11-01-2025 06:45:09” |
| occurence_ongoing | A binary value that lists if an occurrence has been recorded as being chronic or persistent. | True, False |
| reference_source_id | A string that corresponds to the occurrence’s ID in another dataset. Must be matched with ForeignSource. | “ADVR00001” |
| reference_source | A string identifier that corresponds to the data source of the corresponding ForeignID. | “ImmPort” |
| comments | Placeholder for additional comments. |  |

SEA CDM Table:

**Subject**

Data input Template:

<https://github.com/sea-cdm/OSEAN-DB/blob/main/DB-Schema/organism.csv>

Instruction:

<https://sea-cdm.github.io/SEA-CDM/sea-cdm_documentation/organism.html>

| **Property** | **Description** | **Example Usage** |
| --- | --- | --- |
| subject_id | An internal ID that corresponds to an instance of a subject. | “ORGO_1” |
| experiment_id | A foreign key that corresponds to which experiment experimental subject belongs to. | “EXPER_1” |
| group_id | A foreign key that corresponds to which group the experimental subject belongs to; organisms that do not belong to a group are assigned a group value of 1 (to represent a singleton). | “GROUP_1” |
| subject_type | A string representing the general classification of a subject used in an experiment. | “Organism”, “Cell Line” |
| subject_type_id | A string corresponding to the ontology ID for the subject_type attribute. | “OBI_0100026” (organism),  “CLO_0000031” (cell line) |
| species | A string representing the species a subject belongs to; cell lines should list the original species that they belong to. | “Human” |
| species_id | A string representing the species an experimental subject belongs to; this species is identified as equivalent to “OBI_0100026”. | “NCBITaxon:9606” (human) |
| organism_race | A string representing the race or grouping an organism belongs to; this subtype could correspond to a human race or ethnicity. Leave blank for non-humans. | “African American” |
| organism_race_id | A string representing the ontology ID that corresponds to an organism_race. | “NCIT_C16352” (Black or African American) |
| subject_lineage | A string representing the subtype a subject belongs to. | “HeLa Cell”, |
| subject_lineage_id | A string representing the ontology ID that corresponds to an subject_lineage. | “CLO_0003684” (HeLa) |
| organism_age | A double that represents the organism age; this is the lowest age listed for the instance of an experimental subject. This must be paired with an organism_age_unit and organism_age_unit_id. | 46, 57 |
| organsism_age_unit | A string representing the units for an organism_age. | “Day”, “Year” |
| organism_age_unit_id | A string representing the ontological ID used for organism_age_unit. | “UO_0000033” (day),  “UO_0000036” (year), |
| organism_sex | A string representing an organism's biological sex. | “Male”, “Female” |
| organism_sex_id | A string representing the ontological ID used for organism_sex. | “PATO_00000383” (male),  “PATO_00000384” (female) |
| reference_source_id | A string representing the ID listed by the organism in the reference id. |  |
| reference_source | A string representing the organism source for an ontology id | “ImmPort” |
| comments | A string representing comments about the organism class entry. |  |

SEA CDM Table:

**Sample**

Data input Template:

<https://github.com/sea-cdm/OSEAN-DB/blob/main/DB-Schema/sample.csv>

Instruction:

<https://sea-cdm.github.io/SEA-CDM/sea-cdm_documentation/sample.html>

| **Property** | **Description** | **Example Usage** |
| --- | --- | --- |
| sample_id | A primary is used to identify a specific sample. | “SAMP_1” |
| subject_id | A foreign id used to identify a specific subject.  For samples that are composed of multiple organisms, a value of ‘0’ is assigned to subject_id and should be queried via group_id. | “ORGO_1” |
| group_id | The group of subjects that data is collected from. This is done for samples that are generated from multiple subjects. | “GROUP_1”, NULL |
| biosample_collection | A string representing the collection method of the sample. | “Swabbing” |
| biosample_collection_id | A string representing the ontology id for biosample_collection. method of the sample. | “OBI_0002600” (collection of organism from swab) |
| collection_date | A string representing the date the biosample process was done. |  |
| collection_datetime | A string representing the datetime the biosample process was done. |  |
| biosample_type | A string representing the source a biosample is derived from. | “Blood” |
| biosample_type_id | A string representing the ontology id for biosample_type_id. | “OBI_0000655” (blood specimen) |
| biosample_reference_source | A string representing the documentation source for a biosample. | “ImmPort” |
| biopsample_reference_source_id | A string representing the ID for the expsample from the biosample_reference_source. |  |
| expsample_type | A string representing the final processed specimen used for an assay. | “Blood Plasma” |
| expsample_type_id | A string representing the ontological ID listed by expsample type. | “OBI_0100016” (blood plasma specimen) |
| expsample_reference_source | A string representing the documentation source for an expsample. | “GEO” |
| expsample_reference_source_id | A string representing the ID for the expsample from the expsample_reference_source. | GSE_27776 |
| comments | A string used as a placeholder for additional comments or notes. |  |

SEA CDM Table:

**Group**

Data input Template:

<https://github.com/sea-cdm/OSEAN-DB/blob/main/DB-Schema/group.csv>

Instruction:

<https://sea-cdm.github.io/SEA-CDM/sea-cdm_documentation/group.html>

| **Property** | **Description** | **Example Usage** |
| --- | --- | --- |
| group_id | A primary key used to identify a specific group. | “GROUP_1” |
| subject_group | A string representing if the original set of subjects used as a group. | “Organism” |
| subject_group_type_id | A string used to identify the ontology id that corresponds to a subject_group type. | “OBI_0100026” (organisms) |
| sample_group | A string representing the set of samples that are derived from organisms which make up a group. | “Biosample”,  “Expsample” |
| sample_group_type_id | A string used to identify the ontology id that corresponds to a sample_group type. | “OBI_0000671” (sample from organism),  “OBI_0000953” (processed sample) |
| group_size | An integer that stores how many individual organisms are contained within a group. | 36 |
| min_group_age | A double that records the youngest age of a subject in a group. | 17 |
| min_age_unit | A string represent the unit associated with the min_group_age | “Day”, “Year” |
| min_age_unit_id | A string represent the ontology id min_age_unit. | “UO_0000033” (day),  “UO_0000036” (year), |
| max_group_age | A double that records the oldest age of a subject in a group. | 87 |
| max_age_unit | A string represent the unit associated with the max_group_age | “Day”, “Year” |
| max_age_unit_id | A string represent the ontology id max_age_unit. | “UO_0000033” (day),  “UO_0000036” (year), |
| reference_source | A string used to identify the source of the reference_source_id. | “ImmPort” |
| reference_source_id | A string used to identify an external reference for the group. | “ARM512” |
| comments |  |  |

SEA CDM Table:

**Results**

Data input Template:

<https://github.com/sea-cdm/OSEAN-DB/blob/main/DB-Schema/results.csv>

Instruction:

<https://sea-cdm.github.io/SEA-CDM/sea-cdm_documentation/results.html>

| **Property** | **Description** | **Example Usage** |
| --- | --- | --- |
| results_id | An internal ID used to identify assay results in a database. | “RSULT_1” |
| experiment_id | An foreign key used to identify the experiment an assay result is part of. | “EXPER_1” |
| group_id | An ID used to identify the group an assay result is about. Groups that are about an individual organisms or samples are given a special ID. | “GROUP_1” |
| sample_id | A foreign key used to identify the sample a result is about. If used for a group, use “1” as placeholder value. | “SAMP_1” |
| subject_id | A foreign key used to identify the organism results is about. If used for a group, use “1” as placeholder value. | “ORGO_1” |
| documenation_id | An foreign key used to identify the document that contains the documentation for the sort. | “DOCU_1” |
| analysis_type | A string used to identify the current analysis used to generate a result. | Observation, Statistical-Analysis |
| original_assay_type | A string used to identify the original assay type used to generate an assay. This is always an experimental assay. | “DNA Sequence feature detection assay” |
| original_assay_type_id | An ID used to identify the assay ID that an OriginalAssayType is part of. | OBI_0000433 (DNA sequence feature detection assay) |
| datatype | A string used to identify the type of data the results is; either numeric, qualitative, etc. | ‘Image’, ‘Spreadsheet’ |
| datatype_id | An ontology ID that corresponds to the Datatype. | “IAO_0000101” (image),  “SWO_3000001” (spreadsheet format) |
| dataset_size | A string used to identify the size of the results. Currently used as a string due to variable size from matrices or hierarchical data. | “402x27” |
| file_access | A string used to identify the location or web link the results file is located. This is analogous to reference_source in other values. | “https://zenodo.org/records/7407195” |
| file_type | A string used to identify the file type extension the data is stored as (if accessible for computers). | “.csv” |
| replications | An integer that lists how many times an analysis is repeated to generate data; multiple measurements. | 1 |
| comments |  | This uses our VIGET dataset as an example (PMID: 37180100). |

SEA CDM Table:

**Documentation**

Data input Template:

<https://github.com/sea-cdm/OSEAN-DB/blob/main/DB-Schema/documentation.csv>

Instruction:

<https://sea-cdm.github.io/SEA-CDM/sea-cdm_documentation/documentation.html>

| **Property** | **Description** | **Example Usage** |
| --- | --- | --- |
| documentation_id | A primary key used to identify a specific documentation for a document as part of the study. | “DOCU_1” |
| study_id | A foreign key used to identify a study that documentation belongs to. | “STUDY_1” |
| document_name | A string used to represent the name of documentation. | “VIGET: A web portal for study of vaccine-induced host responses based on Reactome pathways and ImmPort data” |
| documentation_type | A string used to represent what the documentation is about; i.e. protocol, paper, results, etc. | “Research Paper” |
| document_type_id | A string used to represent the id of the associated document type. | “IAO_0000312” (publication about investigation) |
| document_file_access | A string representing a URL or file source for a document. | “10.3389/fimmu.2023.1141030” |
| reference_source | A string used to identify the source of documentation that a reference_source_id belongs to. | PubMed |
| reference_source_id | A string used to identify a foreign ID used for a document. This is paired to reference_source . | PMID_1351321 |
| citation | A string representing the citation used for a document. This is used as a shorthand to identify the date of publication. | Brunson T, Sanati N, Huffman A, Masci AM, Zheng J, Cooke MF, Conley P, He Y, Wu G. VIGET: A web portal for study of vaccine-induced host responses based on Reactome pathways and ImmPort data. Front Immunol. 2023 Mar 21;14:1141030. doi: 10.3389/fimmu.2023.1141030. PMID: 37180100; PMCID: PMC10172660. |
| citation_style | A string representing the citation style used for a citation. | “NLM” |
| creator_id | A string used to identify where the role ID is located to identify a creator. | 0000-0000-00000-0000 |
| creator_id_type | A string used to find a unique identifier of a person (if available) or organization who has a role in the creation of the documentation. | “ORCID” |
| creator_role | An OBI ID used to identify the role of a person is involved in, such as creator for documentation. | “reporting agent role”, “responsible party role” |
| creator_role_id | An OBI ID used to identify the role a person is involved in, such as an author or corresponding author, for this document. | OBI_0000068 (reporting agent role),  OBI_0000102 (responsible party role) |

SEA CDM Table:

**Material**

Data input Template:

<https://github.com/sea-cdm/OSEAN-DB/blob/main/DB-Schema/material.csv>

Instruction:

<https://sea-cdm.github.io/SEA-CDM/sea-cdm_documentation/material.html>

| **Property** | **Description** | **Example Usage** |
| --- | --- | --- |
| material_id | A primary key used to identify a specific material. | “MATE_1” |
| material_name | A string used to describe the material name. | “Fluzone” |
| material_name_id | A string used to identify the ontology id that is associated with the material name. | VO_00000064 (Fluzone) |
| organization | A string used to identify the organization responsible for the creation or refinement of a material. | “Pfizer” |
| reference_source_id | A string used to identify an external reference for the organization. | “13-5315170” |
| reference_source | A string used to identify the source of the reference_source_id. | IRS EIN |
| comments |  |  |

SEA CDM Table:

**Ontology**

Data input Template:

<https://github.com/sea-cdm/OSEAN-DB/blob/main/DB-Schema/ontology.csv>

Instruction:

<https://sea-cdm.github.io/SEA-CDM/sea-cdm_documentation/ontology.html>

| **Property** | **Description** | **Example Usage** |
| --- | --- | --- |
| ontology_id | A primary key used internally to identify a specific ontology. | “ONTO_1” |
| documentation_id | An foreign key used to identify the documentation associated with the ontology. | “DOCU_1” |
| ontology_name | A string used to provide the common name that is used as an ontology. | “Ontology of Biomedical Investigations”,  “Ontology of Precision Medical Investigation”,  “Vaccine Ontology” |
| ontology_iri | A string used to identify the ontology IRI that is used for terms within a specific ontology. | “http://purl.obolibrary.org/obo/OBI”  “http://purl.obolibrary.org/obo/OPMI,  “http://purl.obolibrary.org/obo/VO” |
