## Supplemental File 2 for "SEA CDM: Study-Experiment-Assay Common Data Model and Databases for Cross-Domain Data Integration and Analysis": Osean_Viget_ETL_T.py

```
# -*- coding: utf-8 -*-

'''

Created on Mon Mar 24 10:41:46 2025


@author: huffmaar

'''


import pandas as pd

import numpy as np

import sys


if len(sys.argv) == 2:

   VIGET = sys.argv[1]


else:

   try:

       VIGET = sys.argv[2]

       gene_exp = sys.argv[1]   

   except:

       print('Please provide the absolute path of the VIGET metadata file as a command-line argument.')

       sys.exit()


Study_Index = 0


def dummy_loop(term, length):

    #Must be string

    try:

        blanck = []

        print(length)

        for i in range(length):

            blanck.append(length)

            return blanck

    except:

        print('Invalid Inputs')

    return 0


def file_reader(filename):

    try: 

        df = pd.read_csv(filename)

    except:

        print('Could not read as csv file.')

        try:

            df = pd.read_excel(filename)

        except:

            print('Could not read excel file.')

            return 0

    return(df)


def SEA_INIT():

      

    SEA_Study  = pd.DataFrame(columns=['study_id', 'study_name', 'study_description', 'study_type',

                                       'study_type_id', 'study_focus', 'study_focus_id',

                                       'study_keywords', 'study_keyword_ids', 'reference_source_id',

                                       'reference_source', 'comments'])

    

    SEA_Docu = pd.DataFrame(columns=['documentation_id', 'study_id', 'document_name',

                                     'documentation_type', 'documentation_type_id', 'documentation_file_access', 'documentation_reference_source', 'documentation_reference_id', 

                                     'creator_id', 'creator_id_type', 'creator_role', 'creator_role_id', 'citation', 'citation_style','comments'])

    SEA_Exper = pd.DataFrame(columns=['experiment_id', 'study_id', 'documentation_id', 'experiment_type',

                                      'experiment_type_id', 'experiment_control', 'experiment_subject',

                                      'experiment_subject_id', 'reference_source_id', 'reference_source', 'comments'])

    SEA_Inter = pd.DataFrame(columns=['intervention_id', 'experiment_id', 'subject_id',

                                      'material_name', 'material_id', 'dosage', 'dosage_unit', 'dosage_unit_id', 'intervention_route',

                                      'intervention_route_id', 'to_definition', 't0_definition_id', 'intervention_time_unit',

                                      'intervention_time_unit_id', 'reference_source', 'reference_source_id', 'comments'])    

    SEA_Assay  = pd.DataFrame(columns=['assay_id', 'file_access', 'assay_name', 'assay_type', 'assay_type_id', 'reagents',

                                       'device', 'reference_id', 'reference_source', 'comments'])

    SEA_Rslt  = pd.DataFrame(columns=['results_id', 'experiment_id', 'subject_id', 'group_id', 'sample_id', 'documentation_id', 

                                      'assay_id', 'analysis_id','analysis_type', 'original_assay_type',

                                      'original_assay_type_id', 'datatype', 'datatype_id', 'dataset_size', 'file_access', 'file_type', 'replications', 'comments'])

    SEA_Onto  = pd.DataFrame(columns=['ontology_id','documentation_id','ontology_name','ontology_string'])

    SEA_Orgo =  pd.DataFrame(columns=['subject_id', 'experiment_id', 'group_id', 'species',

                                      'species_id', 'organism_lineage', 'organism_lineage_id',

                                      'organism_race', 'organism_race_id', 'organism_age',

                                      'organism_age_unit', 'organism_age_unit_id', 'sex','sex_id', 'reference_id', 'reference_source', 'comments'])

    SEA_Occr  = pd.DataFrame(columns=['occurence_id','subject_id',

                                      'occurence_name', 'occurence_name_id', 'occurence_t0', 'occurence_t0_id', 'reoccurence',

                                      'occurence_severity','occurence_onset','occurence_onset_unit', 'occurence_onset_id',

                                      'occurence_end_time','occurence_end_unit', 'occurence_end_id',

                                      'reference_id', 'reference_source', 'comments' ])

    SEA_Samp  = pd.DataFrame(columns=['sample_id', 'experiment_id', 'subject_id', 'group_id',

                                      'reference_id', 'reference_source', 'biosample_collection', 'biosample_collection_id', 'collection_date', 'collection_datetime',

                                      'collection_time', 'collection_time_unit', 'collection_time_unit_id',

                                      't0_definition', 't0_definition_id', 'biosample_type', 'biosample_type_id', 'biosample_reference_id',

                                      'biosample_reference_source', 'expsample_type',

                                      'expsample_type_id', 'expsample_reference_id',

                                      'expsample_reference_source', 'comments'

])

    SEA_Grop  = pd.DataFrame(columns=['group_id', 'experiment_id', 'subject_group_type',

                                      'subject_group_type_id', 'sample_group_type', 'sample_group_type_id',

                                      'composition', 'group_size', 'min_age', 'min_age_unit',

                                      'min_age_unit_id', 'max_age', 'max_age_unit',

                                      'reference_source_id', 'reference_source', 'comments'])

    SEA_Anal  = pd.DataFrame(columns=['analysis_id', 'group_id', 'documentation_id',

                                                 'input_data', 'input_data_id', 'file_access', 'analysis_name',

                                                 'reference_source_id', 'reference_id', 'comments'])

    SEA_Mats  = pd.DataFrame(columns=['material_id','material_name', 'material_name_id',

                                      'organization', 'reference_source_id', 'reference_source','comments'])

    export = [SEA_Study, SEA_Docu, SEA_Exper, SEA_Inter, SEA_Assay, SEA_Rslt, SEA_Onto, SEA_Orgo, SEA_Occr, SEA_Samp, SEA_Grop, SEA_Mats, SEA_Anal]

    print("All tables initialized.")

    return export    


def read_VIGET(VIGET):

    try: 

        dfVIGET = file_reader(VIGET)

    except:

        print('File could not be found')

        return  0

    return dfVIGET


def load_VIGET(VIGET):

    try:

        VIGET = read_VIGET(VIGET)

    except:

        print('File could not be loaded.')

    template = SEA_INIT()

    STUDY = template[0]

    

    'All files are loaded.'

    return 0


def save_list(df_list):

    filenames = [

        "study.csv", "documentation.csv", "experiment.csv", "intervention.csv",

        "assay.csv", "results.csv", "ontology.csv", "subject.csv",

        "occurrence.csv", "sample.csv", "group.csv", "material.csv", "analysis.csv"

    ]

    

    # These are the tables where .set_index() was used in the code.

    # They MUST be saved with index=True to keep their IDs.

    tables_needing_index = [1, 4, 12] # Documentation, Assay, Analysis

    

    for i, df in enumerate(df_list):

        try:

            filename = filenames[i]

            

            if i in tables_needing_index:

                # Save the Index (because the ID is the index)

                df.to_csv(filename, index=True)

            else:

                # Discard the Index (because the ID is already a column)

                df.to_csv(filename, index=False)

                

            print(f"Successfully saved {filename}")

        except Exception as e:

            print(f"Could not save {filenames[i]}. Error: {e}")

        

    return 0


def indexed(df):

    index_series = df.index.to_series()

    indexed = df

    df['index_id'] = index_series

    return df


def fix_id(df, dfs, ref, refs):

    mapping = ref

    df_replaced = df.replace(mapping)

    return df_replaced


def _source_keys(df1, df2, col1, col2):

    outer_join_df = pd.merge(df1, df2, left_on=col1, right_on=col2, how='outer')


    return outer_join_df


# Intialize SEATEmplates

SEATemplates = SEA_INIT()

viget = read_VIGET(VIGET)


STUDY = SEATemplates[0]

STUDY['reference_source_id'] = viget['immport_study_accession']

STUDY['reference_source'] = 'ImmPort'

STUDY['study_description'] = viget['study_brief_desc']

STUDY['study_name'] = viget['study_title']

STUDY = STUDY.drop_duplicates()

STUDY['study_type'] = 'Clinical Investigation'

STUDY['study_type_id'] = 'OBI_0003697'

STUDY['study_id'] = STUDY.index+1

study_viget = pd.Series({'study_id': 3,

                      'study_name' :'VIGET:Vaccine Immune Gemone Expression Tool',

                      'study_type': 'Hypothesis-Generating Investigation',

                      'study_type_id': 'OBI_0000356',

                      'reference_source_id': '37180100',

                      'reference_source':'PUBMED'})

STUDY = pd.concat([STUDY, study_viget.to_frame().T])


# Early Documentation set up relational identification to repeat for study.

DOCU = SEATemplates[1]


document_list = [

    ['VIGET:Vaccine Immune Gemone Expression Tool', 'Research', '37180100', 'PUBMED', 'https://www.frontiersin.org/journals/immunology/articles/10.3389/fimmu.2023.1141030/full'], 

    ['ImmuneExposureGeneExpression_020922.csv', 'Research Supplement', '37180100', 'PUBMED', 'https://zenodo.org/records/7407195'], 

    ['immport_vaccine_expression_matrix_mapped_merged_approved_genes_091421.csv', 'Research Supplement', '37180100', 'PUBMED', 'https://zenodo.org/records/7407195']]

documentation_ids = 1


citation = 'Brunson T, Sanati N, Huffman A, Masci AM, Zheng J, Cooke MF, Conley P, He Y, Wu G. VIGET: A web portal for study of vaccine-induced host responses based on Reactome pathways and ImmPort data. Front Immunol. 2023 Mar 21;14:1141030. doi: 10.3389/fimmu.2023.1141030. PMID: 37180100; PMCID: PMC10172660.'


for documents in document_list:

    doc_name = documents[0]

    doc_type = documents[1]

    doc_id  = documents[2]

    doc_source = documents[3]

    doc_access = documents[4]

    

    # FIXED: Keys now match the SEA_INIT column names exactly

    document1 =  pd.Series({'documentation_id': documentation_ids,

                              'study_id': 2,

                              'citation':f'{citation}',

                              'citation_style': 'NLM',

                              'document_name': f'{doc_name}',

                              'documentation_type': f'{doc_type}', # Fixed

                              'documentation_reference_id': f'{doc_id}', # Fixed typo

                              'documentation_file_access':f'{doc_access}', # Fixed

                              'documentation_reference_source': f'{doc_source}'}) # Fixed

    DOCU =  pd.concat([DOCU, document1.to_frame().T])

    documentation_ids +=1


SEATemplates[1] = DOCU.set_index('documentation_id')


EXPER =  SEATemplates[2]

EXPER['study_id'] = viget['immport_study_accession']

EXPER['experiment_type'] = 'vaccination response'

EXPER['experiment_type_id'] = 'VO_0000002'

EXPER['experiment_control'] = 0

EXPER['experiment_subject'] = 'human'

EXPER['reference_source'] = 'VIGET'

EXPER['reference_source_id'] = viget['immport_study_accession']+' '+viget['vaccine']

EXPER['file_access'] = 'https://www.frontiersin.org/journals/immunology/articles/10.3389/fimmu.2023.1141030/full'

EXPER = EXPER.drop_duplicates()

EXPER['experiment_id'] = EXPER.index+1

EXPER['study_id'] = EXPER['study_id'].map(STUDY.set_index('reference_source_id')['study_id'])


ONTO = SEATemplates[6]

ontology = pd.Series({'ontology_id': 1,

                           'documentation_id': 'obofoundry.org/ontology/vo',

                           'ontology_name': 'Vaccine_Ontology',

                           'ontology_string': 'VO'})

ONTO = ontology.to_frame().T

SEATemplates[6] = ONTO


import pandas as pd


# Original DataFrame

df = pd.DataFrame({'col1': [1, 2], 'col2': [3, 4]})

print('Original DataFrame:')

print(df)


# New row as a Series

new_row_series = pd.Series({'col1': 5, 'col2': 6})


# Convert Series to a single-row DataFrame

new_row_df = new_row_series.to_frame().T


# Concatenate to add the new row

df_updated = pd.concat([df, new_row_df], ignore_index=True)


print('\nDataFrame with new row added:')

print(df_updated)


GROUP = SEATemplates[10]

GROUP['group_id'] = viget['batch_factor']

GROUP['experiment_id'] = viget['immport_study_accession']+' '+viget['vaccine']

GROUP['sample_group_type_id'] = 'OBI_0000953'

GROUP['sample_group_type'] = 'Expsamples'

GROUP['min_age'] = viget['study_min_age']

GROUP['min_age_unit'] = 'Yr'

GROUP['min_age_unit_id'] = 'UO_0000036'

GROUP['max_age'] = viget['study_max_age']

GROUP['max_age_unit'] = 'Yr'

GROUP['max_age_unit_id'] = 'UO_0000036'

GROUP['reference_source_id'] = viget['batch_factor']

GROUP['reference_source'] = 'VIGET'

GROUP['subject_group_type'] = 'organism'

GROUP['subject_group_type_id'] = 'OBI_0000671'

GROUP = GROUP.drop_duplicates(subset='group_id')


ORGO = SEATemplates[7]

ORGO['reference_source'] = 'ImmPort'

ORGO['species'] = 'human'

ORGO['species_id'] = 'NCBITaxon_9606'

ORGO['group_id'] = viget['immport_study_accession']+' '+viget['batch_factor']

ORGO['sex'] = viget['gender']

ORGO['sex_id'] = 'PATO_0001894'

ORGO['sex_id'][ORGO['sex']=='Male'] = 'PATO_0000384'

ORGO['sex_id'][ORGO['sex']=='Female'] = 'PATO_0000383'

ORGO['experiment_id'] = viget['immport_study_accession']+' '+viget['vaccine']

ORGO['organism_race'] = viget['race']

ORGO['experiment_id'] = ORGO['experiment_id'].map(

    EXPER.set_index('reference_source_id')['experiment_id'])


# FIXED: Changed 'reference_source_id' to 'reference_id' to match the column name

ORGO['reference_id'] = viget['immport_subject_accession'] 

ORGO['reference_source'] = 'ImmPort'

ORGO['age_unit_id'] = 'UO_0000001'


# Now this works because reference_id actually has data

ORGO = ORGO.drop_duplicates(subset = 'reference_id') 

ORGO['subject_id'] = ORGO.index+1


INTER= SEATemplates[3]

INTER['experiment_id'] = viget['immport_study_accession']+' '+viget['vaccine']

INTER['material_name_id'] = viget['immport_immune_exposure_material_id']

INTER['material_name'] = viget['vaccine']

INTER['intervention_time'] = 0

INTER['intervention_unit'] = viget['immport_vaccination_time_unit']

INTER['intervention_time_unit_id'] = 'UO_0000035'

INTER['reference_id'] = 'VIGET'

INTER['T0_definition'] = viget['day_0_def']

INTER['reference_source'] = 'ImmPort'

INTER['subject_id'] = viget['immport_subject_accession']

# keep

INTER['experiment_id'] = INTER['experiment_id'].map(EXPER.set_index('reference_source_id')['experiment_id'])

INTER['subject_id'] = INTER['subject_id'].map(ORGO.set_index('reference_id')['subject_id'])

INTER['intervention_type'] =  'vaccination'

INTER['intervention_type_id'] =  'VO_0000001'

INTER['intervention_route_id'] = 'VO_0000574'

INTER['dosage_unit_id'] = 'OBI_0000984'

INTER = INTER.drop_duplicates(subset='subject_id')

INTER['intervention_id'] = INTER.index+1


SAMP=SEATemplates[9]

SAMP['subject_id'] = viget['immport_subject_accession']

SAMP['group_id'] = viget['batch_factor']

SAMP['experiment_id'] = viget['immport_study_accession']+' '+viget['vaccine']


SAMP['biosample_type'] = viget['biosample_type']

SAMP['expsample_type'] =  viget['type_subtype']

SAMP['expsample_source'] = viget['expsample_repository_name']

SAMP['biosample_reference_id'] = viget['immport_biosample_accession']

SAMP['expsample_reference_id'] = viget['gsm']

SAMP['collection'] = 'blood draw'

SAMP['collection_id'] = 'EFO_0009121'

SAMP['collection_time'] = viget['immport_vaccination_time']

SAMP['collection_time_unit'] = viget['immport_vaccination_time_unit']

SAMP['collection_time_unit_id'] = 'UO_0000035'


#Assignment of additional metadata data.

SAMP['biosample_type_id'] = 'NCIT_C43412'

SAMP['biosample_type_id'][SAMP['biosample_type'] == 'PBMC'] = 'NCIT_C12519'

SAMP['biosample_type_id'][SAMP['biosample_type'] == 'B cell'] = 'CL_0000236'

SAMP['biosample_type_id'][SAMP['biosample_type'] == 'Whole Blood'] = 'NCIT_C12434'


SAMP['group_id'] = SAMP['group_id'].map(GROUP.set_index('reference_source_id')['group_id'])

SAMP['experiment_id'] = SAMP['experiment_id'].map(EXPER.set_index('reference_source_id')['experiment_id'])

SAMP['subject_id'] = SAMP['subject_id'].map(ORGO.set_index('reference_id')['subject_id'])

SAMP['sample_id'] = SAMP.index+1


 

# Assay and Analysis follow this loop.

ASSAY = SEATemplates[4]

ALYS = SEATemplates[12]


zi = 0

for z in viget['gpl'].unique():

    platform = z

    zi += 1

    # FIXED: Mapped 'platform' to 'device' and ensured keys match schema

    assay = pd.Series({'assay_id': zi,

                       'assay_name': 'Blood.RNA-Microarray',

                       'assay_type': 'Experimental Assay',

                       'assay_type_id': 'ECO_0000097',

                       'reagents': 'mRNA',

                       'file_access': '/immport_vaccine_expression_matrix_mapped_merged_approved_genes_091421.csv',

                       'device': f'{platform}'}) # Changed 'platform' to 'device'

    ASSAY = pd.concat([ASSAY, assay.to_frame().T])


# ... (Assay block is above this) ...


# FIXED Analysis Section

alys = pd.Series({'analysis_id': 1,  

                   'analysis_name': 'VIGET LIMMA analysis',

                   # 'group_id': '', # Left empty as in original, or assign if known

                   'documentation_id': 2, # Linked to the Zenodo documentation we created earlier

                   'input_data_id':'.csv',

                   'input_data': 'VIGET',

                   'reference_source': 'GEO / ImmPort',

                   'reference_source_id': 'VIGET', # Matched to schema column

                   'file_access': 'https://zenodo.org/records/7407195/immport_vaccine_expression_matrix_mapped_merged_approved_genes_091421.csv',

                   'comments': '' 

                   })

ALYS = pd.concat([ALYS, alys.to_frame().T])


# Ensure columns line up with the template

SEATemplates[12] = ALYS.set_index('analysis_id')

SEATemplates[4] = ASSAY.set_index('assay_id')


RESULT = SEATemplates[5]


RESULT['sample_id'] = viget['gsm']

RESULT['subject_id'] = viget['immport_subject_accession']

RESULT['group_id'] = viget['batch_factor']

RESULT['experiment_id'] = viget['immport_study_accession']+' '+viget['vaccine']

RESULT['assay_id'] = viget['gpl']

RESULT['datatype'] = 'spreadsheet'

RESULT['datatype_id'] = 'SWO_3000001'

# Information on result categories.

# RESULT['file_size']

RESULT['file_access'] = 'https://zenodo.org/records/7407195/immport_vaccine_expression_matrix_mapped_merged_approved_genes_091421.csv'

RESULT['file_type'] = 'csv'

RESULT['original_assay_type'] = 'Blood.RNA Expression Assay'

RESULT['documentation_id'] = 2

RESULT['replications'] = 1


# REMAP ALL Result_ids

RESULT['experiment_id'] =  RESULT['experiment_id'].map(EXPER.set_index('reference_source_id')['experiment_id'])

RESULT['group_id'] = RESULT['group_id'].map(GROUP.set_index('reference_source_id')['group_id'])

RESULT['subject_id'] = RESULT['subject_id'].map(ORGO.set_index('reference_id')['subject_id'])

RESULT['original_assay_type_id'] = RESULT['assay_id'].map(ASSAY.set_index('device')['assay_id'])

RESULT['results_id'] = RESULT.index+1


MATS = SEATemplates[11]

MATS['reference_id'] = viget['immport_immune_exposure_material_id']

MATS['material_name_id'] = viget['immport_immune_exposure_material_id']

MATS['reference_source'] = 'VIGET'

MATS['material_name'] = viget['vaccine']

MATS = MATS.drop_duplicates(subset='reference_id')

MATS['material_id'] = MATS.index+1


# --- CRITICAL FIX: Update the master list with the modified tables ---

SEATemplates[0] = STUDY

SEATemplates[2] = EXPER

SEATemplates[3] = INTER

SEATemplates[5] = RESULT

SEATemplates[7] = ORGO

SEATemplates[9] = SAMP

SEATemplates[10] = GROUP

SEATemplates[11] = MATS

# ---------------------------------------------------------------------


save_list(SEATemplates)

print('All tables saved.')


unit_test = SEA_INIT()


for unit in range(len(unit_test)):

    #Test for non id columns are matching

    print(len(unit_test[unit].columns), len(SEATemplates[unit].columns))

    if len(unit_test[unit].columns) != len(SEATemplates[unit].columns):

        print(f'Unit test failed on matching columns on table #{unit}')
```
