## Supplemental File 2 for "SEA CDM: Study-Experiment-Assay Common Data Model and Databases for Cross-Domain Data Integration and Analysis": Readme.md

This contains instructions to convert data into VIGET to SEA-CDM format.


Download  the VIGET_metadata file 'immport_vaccine_expression_matrix_mapped_merged_approved_genes_091421.csv',' from Zenodo https://zenodo.org/records/7407195.
Otherwise, you can generate the files using the SEA-CDM_Final.
Use the template files provided for SEA-CDM_template_files and run the SEA-CDM_Final.mwb to load the tables into OSEAN-DB format.

Use the osean_db.schema.mwb to load data into OSEAN-DB.
