## Supplemental File 2 for "SEA CDM: Study-Experiment-Assay Common Data Model and Databases for Cross-Domain Data Integration and Analysis": SEA-CDM.sql

```
-- MySQL Workbench Forward Engineering

SET @OLD_UNIQUE_CHECKS=@@UNIQUE_CHECKS, UNIQUE_CHECKS=0;
SET @OLD_FOREIGN_KEY_CHECKS=@@FOREIGN_KEY_CHECKS, FOREIGN_KEY_CHECKS=0;
SET @OLD_SQL_MODE=@@SQL_MODE, SQL_MODE='ONLY_FULL_GROUP_BY,STRICT_TRANS_TABLES,NO_ZERO_IN_DATE,NO_ZERO_DATE,ERROR_FOR_DIVISION_BY_ZERO,NO_ENGINE_SUBSTITUTION';

-- Schema osean
-- -----------------------------------------------------
DROP SCHEMA IF EXISTS `osean` ;

-- -----------------------------------------------------
-- Schema osean
-- -----------------------------------------------------
CREATE SCHEMA IF NOT EXISTS `osean` DEFAULT CHARACTER SET utf8mb4 ;
SHOW WARNINGS;
USE `osean` ;

SHOW WARNINGS;
-- -----------------------------------------------------
-- Table `osean`.`study`
-- -----------------------------------------------------
DROP TABLE IF EXISTS `osean`.`study` ;

SHOW WARNINGS;
CREATE TABLE IF NOT EXISTS `osean`.`study` (
  `study_id` INT NOT NULL,
  `study_name` VARCHAR(45) NULL,
  `study_description` VARCHAR(450) NULL DEFAULT NULL,
  `study_type` VARCHAR(45) NULL DEFAULT NULL,
  `study_type_id` VARCHAR(15) NULL,
  `study_focus` VARCHAR(45) NULL DEFAULT NULL,
  `study_focus_id` VARCHAR(15) NULL,
  `study_keywords` VARCHAR(450) NULL,
  `study_keyword_ids` VARCHAR(150) NULL,
  `reference_source_id` VARCHAR(45) NULL DEFAULT 'Unknown',
  `reference_source` VARCHAR(45) NULL DEFAULT NULL,
  `comments` VARCHAR(500) NULL,
  PRIMARY KEY (`study_id`))
ENGINE = InnoDB
DEFAULT CHARACTER SET = utf8mb4;

SHOW WARNINGS;

-- -----------------------------------------------------
-- Table `osean`.`experiment`
-- -----------------------------------------------------
DROP TABLE IF EXISTS `osean`.`experiment` ;

SHOW WARNINGS;
CREATE TABLE IF NOT EXISTS `osean`.`experiment` (
  `experiment_id` INT NOT NULL,
  `study_id` INT NULL DEFAULT NULL,
  `documentation_id` INT NULL DEFAULT NULL,
  `experiment_type` VARCHAR(45) NULL DEFAULT NULL,
  `experiment_type_id` VARCHAR(45)  NULL DEFAULT NULL,
  `experiment_control` BINARY NULL,
  `experiment_subject` VARCHAR(45) NULL DEFAULT NULL,
  `experiment_subject_id` VARCHAR(45) NULL DEFAULT "OBO_0000030",
  `reference_source_id` VARCHAR(45) NULL,
  `reference_source` VARCHAR(45) NULL,
  `comments` VARCHAR(255) NULL DEFAULT NULL,
  PRIMARY KEY (`experiment_id`),
  INDEX `studyex` (`study_id` ASC) VISIBLE)
ENGINE = InnoDB
DEFAULT CHARACTER SET = utf8mb4;

SHOW WARNINGS;

-- -----------------------------------------------------
-- Table `osean`.`group`
-- -----------------------------------------------------
DROP TABLE IF EXISTS `osean`.`group` ;

SHOW WARNINGS;
CREATE TABLE IF NOT EXISTS `osean`.`group` (
  `group_id` INT NOT NULL,
  `experiment_id` INT NOT NULL,
  `subject_group_type` VARCHAR(45) NULL DEFAULT NULL,
  `subject_group_type_id` VARCHAR(15) NULL DEFAULT NULL,
  `sample_group_type` VARCHAR(45) NULL DEFAULT NULL,
  `sample_group_type_id` VARCHAR(15) NULL DEFAULT NULL,
  `composition` VARCHAR(45) NULL,
  `group_size` INT NULL,
  `min_age` FLOAT NULL,
  `min_age_unit` VARCHAR(15) NULL,
  `min_age_unit_id` VARCHAR(15) NULL,
  `max_age` FLOAT NULL,
  `max_age_unit` VARCHAR(15) NULL,
  `reference_source_id` VARCHAR(45) NULL DEFAULT NULL,
  `reference_source` VARCHAR(45) NULL DEFAULT NULL,
  `comments` VARCHAR(45) NULL DEFAULT NULL,
  PRIMARY KEY (`group_id`),
  INDEX `experiment_idx` (`experiment_id` ASC) INVISIBLE)
ENGINE = InnoDB
DEFAULT CHARACTER SET = utf8mb4;

SHOW WARNINGS;

-- -----------------------------------------------------
-- Table `osean`.`documentation`
-- -----------------------------------------------------
DROP TABLE IF EXISTS `osean`.`documentation` ;

SHOW WARNINGS;
CREATE TABLE IF NOT EXISTS `osean`.`documentation` (
  `documentation_id` VARCHAR(15) NOT NULL,
  `study_id` INT NULL DEFAULT NULL,
  `document_name` VARCHAR(45) NULL DEFAULT NULL,
  `documentation_type` VARCHAR(45) NULL DEFAULT NULL,
  `documentation_type_id` VARCHAR(15) NULL DEFAULT NULL,
  `documentation_file_access` VARCHAR(450) NULL DEFAULT NULL,
  `documentation_reference_source` VARCHAR(45) NULL DEFAULT NULL,
  `documentation_reference_id` VARCHAR(15) NULL DEFAULT NULL,
  `creator_id` VARCHAR(45) NULL DEFAULT NULL,
  `creator_id_type` VARCHAR(15) NULL DEFAULT NULL,
  `creator_role` VARCHAR(45) NULL DEFAULT NULL,
  `creator_role_id` VARCHAR(15) NULL DEFAULT NULL,
  `citation` VARCHAR(450) NULL DEFAULT NULL,
  `citation_style` VARCHAR(15) NULL DEFAULT NULL,
  `comments` VARCHAR(500) NULL DEFAULT NULL,
  PRIMARY KEY (`documentation_id`),
  INDEX `study_idx` (`study_id` ASC) VISIBLE)
ENGINE = InnoDB
DEFAULT CHARACTER SET = utf8mb4;

SHOW WARNINGS;

-- -----------------------------------------------------
-- Table `osean`.`analysis`
-- -----------------------------------------------------
DROP TABLE IF EXISTS `osean`.`analysis` ;

SHOW WARNINGS;
CREATE TABLE IF NOT EXISTS `osean`.`analysis` (
  `analysis_id` INT NOT NULL,
  `group_id` INT NULL,
  `documentation_id` INT NULL,
  `input_data` VARCHAR(45) NULL,
  `input_data_id` VARCHAR(45) NULL,
  `file_access` VARCHAR(255) NULL,
  `analysis_name` VARCHAR(15) NULL,
  `reference_source_id` VARCHAR(45) NULL,
  `reference_id` VARCHAR(15) NULL,
  `comments` VARCHAR(45) NULL,
  PRIMARY KEY (`analysis_id`),
  INDEX `groupay_idx` (`group_id` ASC) VISIBLE,
  INDEX `documentationay_idx` (`documentation_id` ASC) VISIBLE)
ENGINE = InnoDB;

SHOW WARNINGS;
USE `osean` ;

-- -----------------------------------------------------
-- Table `osean`.`material`
-- -----------------------------------------------------
DROP TABLE IF EXISTS `osean`.`material` ;

SHOW WARNINGS;
CREATE TABLE IF NOT EXISTS `osean`.`material` (
  `material_id` INT NOT NULL,
  `material_name` VARCHAR(45) NULL DEFAULT NULL,
  `material_name_id` VARCHAR(15) NULL DEFAULT 'BFO_0000040',
  `organization` VARCHAR(45) NULL DEFAULT NULL,
  `reference_source_id` VARCHAR(15) NULL,
  `reference_source` VARCHAR(45) NULL,
  `comments` VARCHAR(45) NULL,
  PRIMARY KEY (`material_id`))
ENGINE = InnoDB
DEFAULT CHARACTER SET = utf8mb4;

SHOW WARNINGS;

-- -----------------------------------------------------
-- Table `osean`.`assay`
-- -----------------------------------------------------
DROP TABLE IF EXISTS `osean`.`assay` ;

SHOW WARNINGS;
CREATE TABLE IF NOT EXISTS `osean`.`assay` (
  `assay_id` INT NOT NULL AUTO_INCREMENT,
  `file_access` VARCHAR(255) NULL,
  `assay_name` VARCHAR(45) NULL DEFAULT 'OBI_0000173',
  `assay_type` VARCHAR(45) NULL DEFAULT NULL,
  `assay_type_id` VARCHAR(15) NOT NULL,
  `reagents` LONGTEXT NULL DEFAULT NULL,
  `device` LONGTEXT NULL DEFAULT NULL,
  `reference_id` VARCHAR(20) NULL DEFAULT NULL,
  `reference_source` VARCHAR(50) NULL DEFAULT NULL,
  `comments` VARCHAR(500) NULL,
  PRIMARY KEY (`assay_id`),
  INDEX `reagent_idy_idx` (`device` ASC, `reagents` ASC) VISIBLE,
  INDEX `doccument_idy_idx` (`file_access` ASC) VISIBLE)
ENGINE = InnoDB
DEFAULT CHARACTER SET = utf8mb4;

SHOW WARNINGS;

-- -----------------------------------------------------
-- Table `osean`.`subject`
-- -----------------------------------------------------
DROP TABLE IF EXISTS `osean`.`subject` ;

SHOW WARNINGS;
CREATE TABLE IF NOT EXISTS `osean`.`subject` (
  `subject_id` INT NOT NULL AUTO_INCREMENT,
  `experiment_id` INT NOT NULL,
  `group_id` INT NULL DEFAULT NULL,
  `species` VARCHAR(45) NULL,
  `species_id` VARCHAR(15) NULL,
  `organism_lineage` VARCHAR(45) NULL DEFAULT NULL,
  `organism_lineage_id` VARCHAR(15) NULL DEFAULT NULL,
  `organism_race` VARCHAR(45) NULL,
  `organism_race_id` VARCHAR(15) NULL,
  `organism_age` FLOAT NULL DEFAULT NULL,
  `organism_age_unit` VARCHAR(1) NULL DEFAULT NULL,
  `organism_age_unit_id` VARCHAR(15) NULL,
  `sex` VARCHAR(10) NULL DEFAULT NULL,
  `sex_id` VARCHAR(15) NULL,
  `reference_id` VARCHAR(20) NULL DEFAULT NULL,
  `reference_source` VARCHAR(50) NULL DEFAULT NULL,
  `comments` VARCHAR(450) NULL DEFAULT NULL,
  PRIMARY KEY (`subject_id`),
  INDEX `experiment_idx` (`experiment_id` ASC) VISIBLE,
  INDEX `groups_idx` (`group_id` ASC) VISIBLE)
ENGINE = InnoDB
DEFAULT CHARACTER SET = utf8mb4;

SHOW WARNINGS;

-- -----------------------------------------------------
-- Table `osean`.`intervention`
-- -----------------------------------------------------
DROP TABLE IF EXISTS `osean`.`intervention` ;

SHOW WARNINGS;
CREATE TABLE IF NOT EXISTS `osean`.`intervention` (
  `intervention_id` INT NOT NULL,
  `experiment_id` INT NOT NULL,
  `subject_id` INT NOT NULL,
  `material_name` VARCHAR(100) NULL DEFAULT NULL,
  `material_name_id` VARCHAR(15) NULL DEFAULT 'BFO_0000400',
  `dosage` FLOAT NULL DEFAULT NULL,
  `dosage_unit` VARCHAR(45) NULL DEFAULT NULL,
  `dosage_unit_id` VARCHAR(15) NULL DEFAULT NULL,
  `intervention_route` VARCHAR(50) NULL DEFAULT 'ENVO_0000001',
  `intervention_route_id` VARCHAR(15) NULL DEFAULT NULL,
  `t0_definition` VARCHAR(50) NULL DEFAULT NULL,
  `t0_definition_id` VARCHAR(15) NULL,
  `intervention_time_unit` FLOAT NULL,
  `intervention_time_unit_id` VARCHAR(15) NULL,
  `reference_source` VARCHAR(45) NULL,
  `reference_source_id` VARCHAR(45) NULL,
  `comments` VARCHAR(500) NULL,
  INDEX `experiment_idx` (`experiment_id` ASC) VISIBLE,
  INDEX `organismint_idx` (`subject_id` ASC) VISIBLE)
ENGINE = InnoDB
DEFAULT CHARACTER SET = utf8mb4;

SHOW WARNINGS;

-- -----------------------------------------------------
-- Table `osean`.`occurence`
-- -----------------------------------------------------
DROP TABLE IF EXISTS `osean`.`occurence` ;

SHOW WARNINGS;
CREATE TABLE IF NOT EXISTS `osean`.`occurence` (
  `occurence_id` INT NOT NULL,
  `subject_id` INT NOT NULL,
  `occurence_name` VARCHAR(45) NULL DEFAULT NULL,
  `occurence_name_id` VARCHAR(15) NULL,
  `occurence_severity` INT NULL DEFAULT NULL,
  `occurence_onset_date` DATE NULL DEFAULT NULL,
  `occurence_onset_datetime` DATETIME NULL DEFAULT NULL,
  `occurence_end_date` DATE NULL DEFAULT NULL,
  `occurence_end_datetime` DATETIME NULL DEFAULT NULL,
  `occurence_ongoing` BINARY NULL, 
  `reference_source` VARCHAR(45) NULL,
  `reference_source_id` VARCHAR(45) NULL,
  `comments` VARCHAR(500) NULL,
  PRIMARY KEY (`occurence_id`),
  INDEX `subject_id` (`subject_id` ASC) VISIBLE)
ENGINE = InnoDB
DEFAULT CHARACTER SET = utf8mb4;

SHOW WARNINGS;

-- -----------------------------------------------------
-- Table `osean`.`ontology`
-- -----------------------------------------------------
DROP TABLE IF EXISTS `osean`.`ontology` ;

SHOW WARNINGS;
CREATE TABLE IF NOT EXISTS `osean`.`ontology` (
  `ontology_id` INT NOT NULL,
  `documentation_id` VARCHAR(15) NULL DEFAULT NULL,
  `ontology_name` VARCHAR(45) NULL DEFAULT 'BFO_0000040',
  `ontology_iri` VARCHAR(45) NULL DEFAULT NULL,
  PRIMARY KEY (`ontology_id`),
  INDEX (`documentation_id` ASC) VISIBLE)
ENGINE = InnoDB
DEFAULT CHARACTER SET = utf8mb4;

SHOW WARNINGS;

-- -----------------------------------------------------
-- Table `osean`.`sample`
-- -----------------------------------------------------
DROP TABLE IF EXISTS `osean`.`sample` ;

SHOW WARNINGS;
CREATE TABLE IF NOT EXISTS `osean`.`sample` (
  `sample_id` INT NOT NULL,
  `experiment_id` INT NOT NULL,
  `subject_id` INT NOT NULL DEFAULT '-1',
  `group_id` INT NOT NULL DEFAULT 0,
  `reference_id` VARCHAR(45) NULL DEFAULT NULL,
  `reference_source` VARCHAR(45) NULL DEFAULT NULL,
  `biosample_collection` VARCHAR(45) NULL,
  `biosample_collection_id` VARCHAR(15) NULL,
  `collection_date` DATE NULL,
  `collection_datetime` DATETIME NULL,
  `collection_time_unit_id` VARCHAR(15) NULL,
  `to_definition` VARCHAR(45) NULL,
  `to_definition_id` VARCHAR(15) NULL,
  `biosample_type` VARCHAR(45) NULL,
  `biosample_type_id` VARCHAR(45) NULL,
  `biosample_reference_id` VARCHAR(45) NULL,
  `biosample_reference_source` VARCHAR(45) NULL,
  `expsample_type` VARCHAR(45) NULL,
  `expsample_type_id` VARCHAR(45) NULL,
  `expsample_reference_id` VARCHAR(45) NULL,
  `expsample_reference_source` VARCHAR(45) NULL,
  `comments` VARCHAR(450) NULL,
  PRIMARY KEY (`sample_id`),
  INDEX `group_idx` (`group_id` ASC) VISIBLE,
  INDEX `experiment_idx` (`experiment_id` ASC) VISIBLE,
  INDEX `subject_idx` (`subject_id` ASC) VISIBLE)
ENGINE = InnoDB
DEFAULT CHARACTER SET = utf8mb4;

SHOW WARNINGS;

-- -----------------------------------------------------
-- Table `osean`.`results`
-- -----------------------------------------------------
DROP TABLE IF EXISTS `osean`.`results` ;

SHOW WARNINGS;
CREATE TABLE IF NOT EXISTS `osean`.`results` (
  `results_id` INT NOT NULL AUTO_INCREMENT,
  `experiment_id` INT NULL,
  `subject_id` INT NULL,
  `group_id` INT NULL DEFAULT NULL,
  `sample_id` INT NULL,
  `documentation_id` INT NULL,
  `assay_id` VARCHAR(15)  NULL DEFAULT NULL,
  `analysis_id` INT NULL,
  `analysis_type` VARCHAR(45) NULL DEFAULT NULL,
  `original_assay_type` VARCHAR(45) NULL DEFAULT NULL,
  `original_assay_type_id` VARCHAR(20) NULL DEFAULT NULL,
  `datatype` VARCHAR(45) NULL,
  `datatype_id` VARCHAR(45) NULL,
  `dataset_size` VARCHAR(50) NULL DEFAULT NULL,
  `file_access` VARCHAR(255) NULL DEFAULT NULL COMMENT '0 means data structure can be reordered without data loss.\\n1 means data stucture is necessarily ordered with their variables.',
  `file_type` VARCHAR(15) NULL,
  `replications` INT NULL,
  `comments` VARCHAR(500) NULL,
  PRIMARY KEY (`results_id`),
  INDEX `experiment_idx` (`experiment_id` ASC) VISIBLE,
  INDEX `assayar_idx` (`assay_id` ASC) VISIBLE,
  INDEX `samplear_idx` (`sample_id` ASC) VISIBLE,
  INDEX `documensar_idx` (`documentation_id` ASC) VISIBLE,
  INDEX `analysisar_idx` (`analysis_id` ASC) VISIBLE,
  INDEX `groupar_idx` (`group_id` ASC) VISIBLE,
  INDEX `organismar_idx` (`subject_id` ASC) VISIBLE)
ENGINE = InnoDB
DEFAULT CHARACTER SET = utf8mb4;

SHOW WARNINGS;

SET SQL_MODE=@OLD_SQL_MODE;
SET FOREIGN_KEY_CHECKS=@OLD_FOREIGN_KEY_CHECKS;
SET UNIQUE_CHECKS=@OLD_UNIQUE_CHECKS;
```
