## Supplemental Tables for "SEA CDM: Study-Experiment-Assay Common Data Model and Databases for Cross-Domain Data Integration and Analysis"

**Supplemental Tables of File.**

**Supplemental Table 1. General metadata tables and columns in OSEAN DB.** Each column indicates a variable or metadata ids. All variables are represented via a name and matched ontology ID. All metadata columns after the semi-colon are metadata ids used by OSEAN DB internally (for foreign key) or externally (to reference sources).

| **SEA Classes** | **Representative Metadata Columns** |
| --- | --- |
| Study | study_type, study_focus, study_keywords;  study_name, study_description, reference_source_id, reference_source, comments |
| Documentation | documenation_type, documentation_file_access, documentation_reference_source, document_citation, documentation_citation_style, creator_id, creator_id_type, creator_role;  study_id, document_name, documentation_reference_id, documentation_reference_source, comments |
| Experiment | experiment_type, experiment_control, experiment_subject;  study_id, documentation_id, reference_source_id, reference_source, comments |
| Assay | assay_type, organism_inclusion, reagents, platform; file_access, reference_source_id, reference_source, documentation_id |
| Result | datatype, original_assay_type, filetype;  analysis_id, documentation_id, sample_id, group_id |
| Analysis | analysis_type, analysis_input, analysis_data, parameters, hyperparameters, reference_id, reference_source; group_id, experiment_id, reference_source, reference_source_id |
| Subject | age, sex, subject_type, species, race, ethnicity, strain, reference_source_id, reference_source; group_id, experiment_id |
| Groups | subject_group_type, sample_group_type, composition, group_size, min_age, min_age_unit, max_age, max_age_unit, reference_source_id, reference_source; experiment_id |
| Intervention | material, intervention_time, dose, route, intervention_type; experiment_id, subject_id |
| Sample | collection, collection_time, biosample_source, experimental_sample_type; experiment_id, subject_id, group_id |
| Occurrence | occurrence, occurence_type, occurence_severity, occurence_onset, occurence_end; subject_id |
| Ontology | ontology_name, ontology_IRI, ontology_code; documentation_id |
| Material | material_name, material_type, supplier |

**Supplemental Table 2. Conversion of VIGET Metadata into OSEAN Format.** Each box represents a table containing data related to VIGET. Class entries are listed by class.row format. Related SEA-CDM class entries represent additional annotations done to VIGET. One row, expsample_to_multiple_gsm_flag can be answered via querying the data in OSEAN and thus dropped.

| **VIGET Metadata name** | **SEA CDM Class Entry** | **Related SEA CDM Class Entries** |
| --- | --- | --- |
| immport_subject | subject.reference_source_id | organism.reference_source("ImmPort") |
| gender | subject.organism_sex | organism.species("Human") AND organism.species_ID(NCBITaxon:91056) |
| race | subject.organism_race |  |
| Immport_immune_exposure_material_id | intervention.material_id | ontology.ontology(“Vaccine Ontology”) |
| vaccine | intervention.material_name  material.material_name | material.material_name AND intervention.material_name_id("VO_0000001") |
| biosample_id | sample.biosample_id | sample.biosample_reference_source("ImmPort") |
| immport_vaccination_time | intervention.intervention_time |  |
| immport_vaccination_time_unit | intervention.intervention_time_unit | intervention.time_unit_id |
| day_0_def | intervention.t0_definition |  |
| biosample_type | sample.biosample_type | sample.collection_type(“Blood”) |
| immport_biosample_accession | sample.biosample_reference_id | sample.biosample_repository_source("ImmPort) |
| expsample_repository_name | sample.expsample_reference_source_id(“GEO”) |  |
| gsm | sample.expsample_reference_id |  |
| expsample_to_multiple_gsm_flag |  |  |
| GSE | experiment.experiment_id | results.foreign_repository("GEO") AND assay_results.file_type("Spreadsheet")  results.original_assay(“RNAExpression.Assay”)AND results.data_respository_name("ImmPort") |
| GPL | assay.platform |  |
| platform_desc |  |  |
| batch_factor | group.reference_source_id | group.reference_source(“VIGET”) |
| type_subtype | assay.material AND  sample.biosample_source |  |
| viget_csv_file | results.documenation_link | results.reference_id("GEO") AND results.file_format("VIGET")  results.original_assay_type(“Gene Expression Assay”) |
| age_group | group.min_age AND group.max_age |  |
| study_min_age | group.min_age | group.min_age_unit(“year”) |
| study_max_age | group.max_age | group.max_age_unit(“year”) |
| study_brief_desc | study.description |  |
| study_tile | study.title |  |
| subtype | sample.exp_sample_type |  |

**Supplemental Table 3. Mapping of common ImmPort Core Tables to SEA-CDM Classes.**

| **ImmPort Tables** | **SEA CDM Classes** |
| --- | --- |
| Study | Study |
| Study Condition or Disease | Study, Occurence |
| Inclusion/Exclusion | Study |
| Study-Linked Documentation (Study File, Study Link, Study PubMed, Study Personnel) | Documentation |
| Arm or Cohort | Group |
| Subject | Subject |
| Adverse Event | Occurence |
| Experiment | Experiment |
| Planned Visit | Experiment |
| Protocol | Documentation |
| Immune Exposure | Intervention |
| Intervention | Intervention |
| Treatment | Intervention |
| Reagent | Material |
| Assessment Panel | Analysis, Result |
| Biosample | Sample |
| Control Sample | Sample |
| ExpSample | Sample |
| Lab Test Panel | Analysis, Result |
| Assessment Component | Analysis, Assay, Result |
| Lab Test | Assay, Result |
| ExpSample Results (ELISA, ELISPOT, Flow Cytometry, Mass cytometry, Gene Expression, Genotyping, HAI Result,  HLA typing Result, KIR Typing Result, Lumines/MBAA, Metabolomics, Neutralizing Antibody Tier, Proteomics, Other) | Assay, Result |
| Ontologies (Vaccine Ontology, Disease Ontology, Human Phenotype Ontology, Uberon Anatomy Ontology, Clinical Measurement Ontology, Cell Ontology, Protein Ontology, Gene Ontology, NCBI-Taxonomy, Ontology Biomedical Intervention) | Ontology |

**Supplemental Table 4. Mapping of common CellxGene Metadata Variables Category to SEA CDM categories**.

| **CellxGene Metadata Name** | **SEA CDM** |
| --- | --- |
| Assay | assay.assay_type |
| Cell Type | sample.expsample_type AND  subject.subject_type(“cell line”) |
| Developmental Stage | subject.age AND subject.age_unit |
| Disease | subject.name AND subject.type(“disease”) |
| Sex | subject.sex |
| Self_Reported_Ethnicity | subject.race |
| Species | subject.species |
| Suspension | sample.expsample_type |
| PatientID | subject.reference_id |
| Tissue | sample.biosample_source |
